# Eulerian–Lagrangian and thin film modeling of drug delivery under physiologically varying breathing in a realistic lung model

**DOI:** 10.1101/2025.07.01.662688

**Authors:** Sameer Kumar Verma, Saurabh Bhardwaj, Kishore Singh Patel, B. Kiran Naik

**Affiliations:** Department of Mechanical Engineering, National Institute of Technology Rourkela, Odisha-769008; Sustainable Thermal Energy Systems Laboratory (STESL), Department of Mechanical Engineering, National Institute of Technology Rourkela, Rourkela, Odisha - 769008; Department of Applied Mechanics, Motilal Nehru National Institute of Technology Allahabad, Prayagraj-211004

**Keywords:** Respiratory drug delivery, CT-scan model, Discrete phase model, Eulerian wall film model, COPD breathing, CFD

## Abstract

This work presents a computational approach to investigate antiviral aerosol deposition in anatomically realistic human airways under various physiological inhalation conditions. Antiviral drugs are represented as fine liquid droplets that form a thin coating on airway surfaces, primarily targeting early infection sites. The suggested method combines an Eulerian-Lagrangian discrete phase model (DPM) to track droplet motion and an Eulerian wall film (EWF) model to mimic the evolution of deposited liquid films. This combination effectively addresses the shortcomings of previous studies that employed DPM exclusively, which ignored the post-deposition redistribution dynamics of liquid drugs on airway surfaces. Furthermore, the effects of the distressed breathing of patients with chronic obstructive pulmonary disease (COPD) are compared with both idealized and physiologically realistic breathing patterns in healthy individuals. The study assesses airflow parameters, film thickness, deposition efficiency, and surface area coverage for aerosol particles from 1 to 10 µm. Results reveal that distressed breathing patterns of COPD patients significantly alter the deposition preferences between the upper (~5.28%) and lower lobes (~2.52%) compared to the equivalent ideal and realistic healthy inhalations. Moreover, the distressed breathing also limits the drug penetration into deeper generations, as the highest surface coverage is observed at the carinal region rather than the usual left lower lobe (LLL) found in ideal and realistic healthy breathing cases. Such deposition contrasts highlight the importance of optimizing inhalation therapy and device designs for individuals with obstructive airway diseases.

## 1. Introduction

Understanding the dynamics of respiration is critical for determining its role in human health and disease. Among its various functions, the respiratory system serves as a primary interface for airborne particles entering the body. Once inhaled, these particles can accumulate on the airway surfaces, affecting respiratory health. This deposition process is particularly concerning because it may allow harmful contaminants to enter the lungs, potentially causing pulmonary and cardiovascular diseases (Kelly & Fussell, 2011). However, the respiratory system can also be used to transport medicinal substances. Inhaled drug particles are frequently employed in treating respiratory disorders, providing a direct and efficient method of administering drugs (Dhanani et al., 2016; Dolovich et al., 2019). Inhalation therapy is an efficient technique of respiratory drug delivery to cure various respiratory ailments, such as asthma, chronic obstructive pulmonary disease (COPD), etc. Fig. 1(a) depicts the schematics of the major diseases caused by respiratory viruses and their infection mechanisms. The efficacy of drug delivery to specific airway sites is widely influenced by a range of factors, such as the shape and size of drug particles, the physical conditions of the patient, and the choice of inhalation devices such as pressurized metered dose inhalers (pMDIs), nebulizers, and dry powder inhalers (DPIs) (Fig. 1(b)) (Christou et al., 2021; Dhanani et al., 2016).

**Fig. 1.**
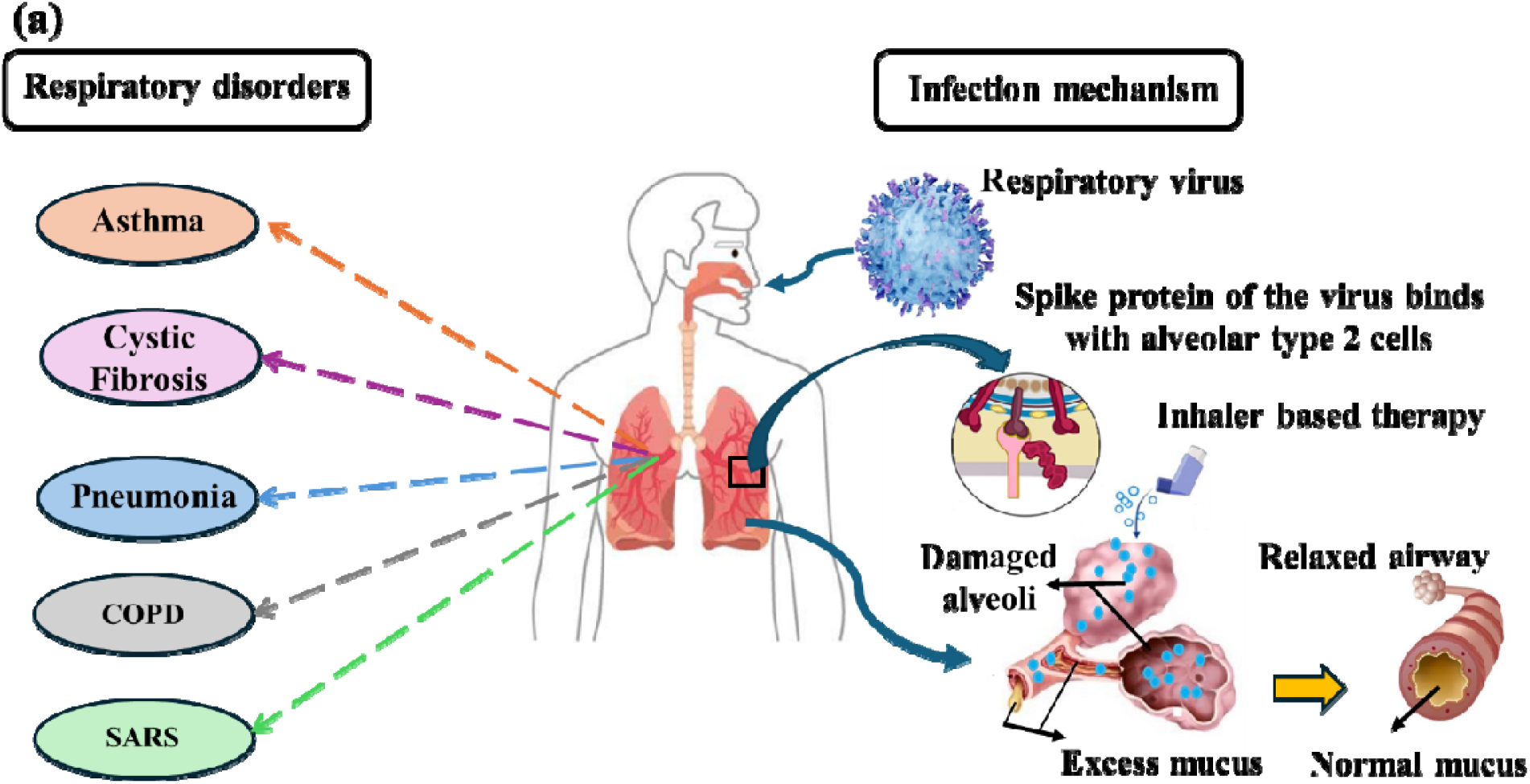
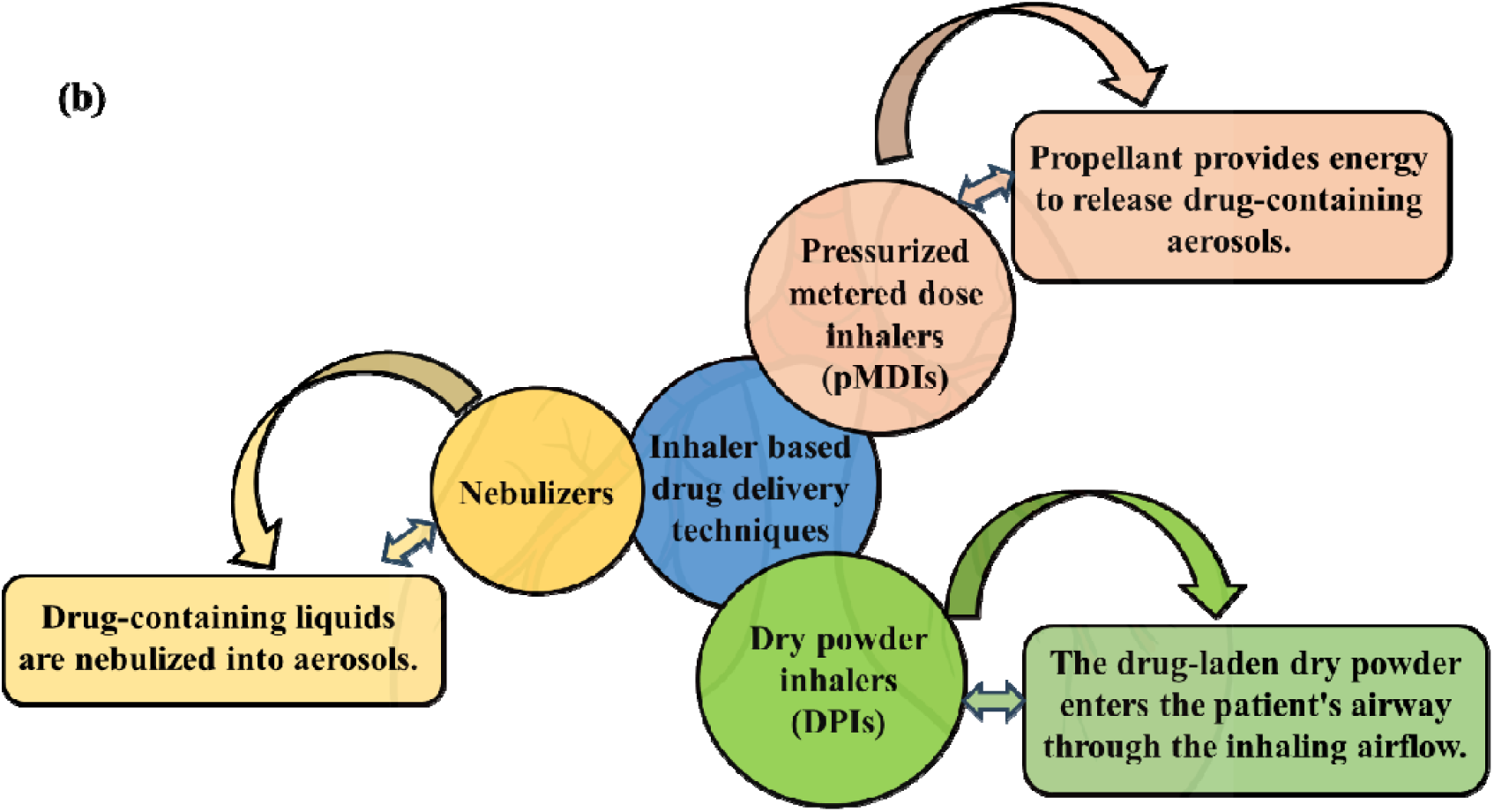
Schematic representations to illustrate (a) the respiratory disorders and the infection mechanism of the respiratory virus that results in airway inflammation, alveolar damage, and increased mucus production, and (b) different inhaler-based direct drug delivery approaches effective in neutralizing the pulmonary viruses.

Numerous experimental research and computational fluid dynamics (CFD) simulations have been carried out to predict the trajectory and deposition of drug particles in the human lung (Ahookhosh et al., 2019; Farghadan et al., 2019; Prinz et al., 2025; Sadeghi et al., 2025; Wu et al., 2022). Weibel’s geometry, often known as the ideal and symmetric lung geometry (Weibel, 1963), helps understand particle trajectory and deposition. Xu & Yu (1986) performed a theoretical investigation based on Weibel’s model for up to generation 23 (G23) to evaluate the effect of age on aerosol deposition for various particle sizes. Comer et al. (2000) used Weibel’s model to replicate particle deposition patterns between G3 and G5. Using a simplified human respiratory tract model, Rahman et al. (2021) investigated the depositions of 5, 10, and 20 μm particles for people of 50, 60, and 70-year age groups. They reduced the airway diameter by 10% for each 10 years of age rise. The study discovered that aging affects deposition efficiency, with particles accumulating more in the upper respiratory tracts in 70-year-olds than in 50-year-olds’ lungs, and the escape rate of the particles decreases with age. Moreover, Rahman et al. (2022) also investigated the deposition of nano-sized particles of 5, 10, and 20 nm up to G11 using a simple lung airway model. They discovered that 5 nm particles had much greater deposition efficiency in each generation than the 10 and 20 nm particles. Verma et al. (2024) have done a computational analysis of Weibel’s geometry (G0-G4) to calculate the more realistic particle deposition by employing the discrete phase model (DPM) and Eulerian wall film (EWF) approach. The above literature demonstrates that CFD is a robust tool for estimating the airflow and particle dynamics inside the respiratory tract.

Further studies have extended the analysis of particle deposition dynamics to patient-specific human lung geometries, offering more profound insights into airflow behavior and deposition patterns in the airways (Bhardwaj et al., 2022; M. M. Rahman et al., 2024; Samanta et al., 2025). Oakes et al. (2018) examined airflow across various developmental stages, adults, children, and infants, to better understand age-dependent flow dynamics. Beni et al. (2021) used a realistic lung model to analyze severe acute respiratory syndrome (SARS) virus deposition patterns under varying constant flow rates and aerosol sizes, although their study was confined to the nasal cavity and trachea without extending into the deeper lung generations. Taheri et al. (2021) reconstructed a human airways model from computed tomography (CT) scan to explore how swirling inlet conditions and different particle release strategies affect deposition efficiency. Wedel et al. (2022) compared steady-state simulation results with transient shear stress transport (SST) k-ω and detached eddy simulation (DES) models to evaluate particle deposition behavior in CT scan-based male and female lung geometries. The study considered various particle size distributions representing different respiratory activities, such as coughing, sneezing, and normal breathing. Islam et al. (2021) analyzed the transport and deposition of SARS-CoV-2 virus particles in a CT scan G17 lung model, focusing on their movement toward the terminal airways.

The investigations mentioned above applied a constant breathing rate as an inlet condition. However, during drug inhalation, the rate at which airflow reaches the lungs through the mouth varies in a cyclical pattern rather than remaining constant. Some studies have considered transient airflow in the human lung (Atzeni et al., 2021; Farnoud et al., 2020; Hamza et al., 2025; Jedelsky et al., 2012; Lambert et al., 2011; Vulović et al., 2022). Paul et al. (2021) replicated the deposition of cigarette smoke particles in patient-specific human lung airways extending up to the G6 of airway bifurcations, including a realistic inhalation profile. Naseri et al. (2017) employed a sinusoidal airflow rate to investigate the path of microparticles traveling from the nostrils to the trachea. The study examined two different inhalation conditions, rest and moderate, allowing for analysis of how varying respiratory patterns influence particle transport. Makris et al. (2014) adopted a sinusoidal airflow rate with two frequencies normal and high, to model airflow and particle deposition between generations G3 and G4. Tiwari et al. (2021) employed a transient inhalation profile through a DPI to predict particle deposition in the G6 lung airways model. Bhardwaj et al. (2022) computationally examined how the transient motion of glottal structures affects inhalation therapy using a realistic airway geometry under a DPI inhalation profile. Their results revealed that glottal deformation in the conducting airways significantly influences the localized deposition of smaller micron-sized particles. Grgic et al. (2006) displayed that deposition efficiency changes between transient and continuous flow patterns. They employed an escalating flow of 45 liters per minute (LPM), followed by a rapid deceleration to zero.

Limited studies are available on disease-specific respiratory drug delivery. These studies are discussed here, explicitly considering their direct relevance to the current work. In this direction, the focus of the early studies (Farkas et al., 2024; Horváth et al., 2017, 2020; Tajiri et al., 2022) was not precisely the analysis of drug delivery efficacy but establishing the breathing patterns of the COPD and asthma patients. The subsequent work by Vitacca et al. (1999) and Colasanti et al. (2004) discovered that the COPD patients recovering from an exacerbation have a contrastingly different breathing pattern than healthy individuals. The characteristics of COPD are often described by tidal breathing, which suggests short, forceful, and rapid inhalation, unlike that in healthy persons. In recent studies, Farkas et al. (2020) numerically and experimentally compared the deposition efficiencies of mannitol and chitosan particles for severe asthmatic and healthy breathing cases. The comparison between the numerical and experimental studies shows good agreement on the ideal and realistic geometries. However, the ideal geometry they consider is only taken up to G2, starting from the trachea. Williams et al. (2022) assessed particle deposition in three anatomical airway models, depicting healthy, lung cancer, and cystic fibrosis states, and compared deposition outcomes using two breathing profiles, one representing a healthy person and the other a COPD patient. Their findings were primarily concerned with overall deposition efficiency, with little emphasis on how impaired breathing patterns affect deposition across specific lung zones. Furthermore, they applied the COPD breathing profile to a different lung disease model, known as cystic fibrosis. As per Colasanti et al. (2004), the COPD and cystic fibrosis diseases had separate inhaling patterns, implying that such an overlap may not effectively represent physiological conditions. Understanding how regional and local deposition responds to different inhalation profiles would enable clinicians to develop a tailor-made inhalation therapy based on the patient’s conditions. Moreover, it would support the development of smart inhalers to achieve drug delivery at target sites within the respiratory tract.

### 1.1. Literature Gap and Novelty of the Study

#### 1.1.1. Shortcomings of existing numerical techniques

The precise prediction of drug particle deposition inside the human respiratory tract is crucial in understanding the therapeutic efficacy of the drug delivery. Among the various numerical techniques, the DPM is commonly used by most researchers to simulate drug particle transport in the human airways. This method can largely track the generic flow feature of the bulk air medium, deposition efficiency of the particles, and their escape rate by a combined Eulerian-Lagrangian approach. The DPM approach can track the particle deposition parameters over a specified zone in an averaged sense rather than their exact pinpointed locations within the zone itself. Moreover, it generally assumes that once a particle touches (regardless of its inertia and strike angle) a surface, it remains stationary under a trapped boundary condition (Rahman et al., 2022; Tiwari et al., 2021) without undergoing any additional interaction. This ignores several post-deposition events, such as:

*(a) Post-impaction dynamics:* Although particle deposition modeling in the respiratory system has advanced significantly, most studies concentrate on the point of impact, often treating particle-wall interaction as a final outcome. In many inhaler-based drug delivery techniques, e.g., pMDI and nebulizers, the antiviral drugs are inhaled in the form of liquid particles or droplets. The trapped boundary condition used in DPM ignores critical phenomena such as the redistribution of deposited liquid, potential detachment under airflow-induced shear, or the formation of secondary droplets. The absence of such a post-impaction study limits our understanding of how deposited particles behave over time and how this affects therapeutic delivery and retention in the lungs.
*(b) Liquid film spread and area coverage:* Inhaled liquid formulations, particularly those from devices such as pMDIs, generate a thin liquid layer of particles when deposited. However, most existing computational models do not address how this liquid coating spreads across the airway surfaces. As a result, they provide only an approximate view of surface coverage, which is typically represented in terms of average deposition across a wider area of the specified zone. This prevents the identification of surface area coverage and film thickness, which are critical in assessing drug delivery efficiency. Furthermore, differences in film thickness and the impact of airflow in dispersing deposited particles remain untapped. As a result, there is a pressing need for models that can capture the dynamic evolution of surface films in order to estimate localized drug distribution better.

#### 1.1.2. Disease-specific investigations

Despite the increasing utilization of advanced computational models in pulmonary drug delivery, a significant gap remains in the comprehensive comparison of ideal, healthy, and disease-specific breathing patterns, particularly with the stiff breathing generally encountered in COPD and asthmatic cases. This comparison is significant in the recalibrations and subtle modifications in the existing inhalation therapy, as the default inhalers are primarily designed according to ideal or healthy breathing conditions. Although a few of the previous studies have compared healthy and disease-specific breathing profiles, a limited understanding of zone-wise or lobe-wise depositions is available for the impaired inhalation profile. Moreover, the effects of detailed particle size variations in zonal depositions are also not fully understood. The details of disease-specific studies are summarized in a paragraph above Section A. An extensive understanding of zone-wise deposition is crucial in delivering a precise dose of medication to the targeted site to ensure better efficacy. Patients with COPD exhibit short, forceful, and rapid inhalation cycles, which significantly alter airflow dynamics and modify the particle transport and deposition patterns. Therefore, there is a need for integrated modeling approaches that account for realistic breathing waveforms across various medical conditions and inhaled particle sizes to improve drug delivery prediction accuracy.

#### 1.1.3. Novelty and clinical significance of the work

To bridge the above gaps, this study presents a novel computational approach by coupling the EWF model with the Lagrangian DPM to overcome the existing limitations in respiratory drug delivery. The proposed method precisely tracks droplet trajectories and resolves the post-deposition dynamics, such as localized surface coverage, variations in film thickness, and the shear-induced redistribution of the thin layer due to surrounding airflow. The EWF approximations model the film dynamics along the radial directions. Therefore, this method serves as an alternative to conventional interface capturing methods (e.g., volume of fluid, level set, phase field, etc.), which demand a very fine mesh near the wall to resolve thin film dynamics, resulting in high computational costs. Besides the thin film tracking, coupling DPM with the Eulerian approximations also enables us to track the secondary particles released from this film and reatomize them due to splashing, stripping, and separation processes.

Adding further novelty, the study also examines the influence of disease-specific breathing patterns, particularly those observed in individuals with COPD, compared to ideal and healthy breathing conditions. Using a CT scan-based human airway geometry extending up to the G7, the simulations evaluate how such pathological breathing patterns influence drug delivery outcomes. The simulation results indicate that distressed breathing significantly alters the deposition dynamics, resulting in higher deposits along the upper lobes. Moreover, the precise deposition information captured using the film thickness parameter indicates higher localized deposition along the carina region. The current results suggest an unusual deposition pattern compared to previously reported findings. Clinically accurate prediction of film spread and surface coverage with disease-specific inhalation is critical in effective drug delivery. These characteristics define the therapeutic efficacy by ensuring that the drug reaches the target site uniformly. Failing to address them can result in an inadequate dose and reduced therapeutic benefit. Thus, this work signifies the importance of incorporating disease-specific breathing dynamics to enhance the predictive accuracy of pulmonary drug delivery models using an advanced modeling framework.

## 2. Methodology

The mechanisms that cause particle deposition in the respiratory tract are impaction, diffusion, interception, and sedimentation (Choi & Kim, 2007; Hofmann, 2011). The effectiveness of each mechanism is influenced by factors such as airway geometry, particle properties (e.g., size, shape, and density), and the breathing rate. The numerical method consists of two sections: an airflow model that simulates airflow within the human airways and a particle transport model that predicts particle movement based on airflow velocity. The present analysis is performed using ANSYS FLUENT software (2023 R1).

### 2.1. Respiratory tract model and breathing maneuver

This study presents a three-dimensional (3D) anatomical lung model of an adult male created using CT scan data. In this investigation, we employ the SimInhale benchmark anatomical model of the respiratory system, which was initially developed by Lizal *et al*.(Lizal et al., 2012, 2015) and used further by Koullapis *et al*.(P. G. Koullapis et al., 2018) and Bhardwaj *et al*.(Bhardwaj et al., 2022) The tracheobronchial model spans from the trachea to the G7 with different lobes as the right lobe (RL), right upper lobe (RUL), right lower lobe (RLL), left lobe (LL), left upper lobe (LUL), and left lower lobe (LLL), as shown in Fig. 2(a).

**Fig. 2.**
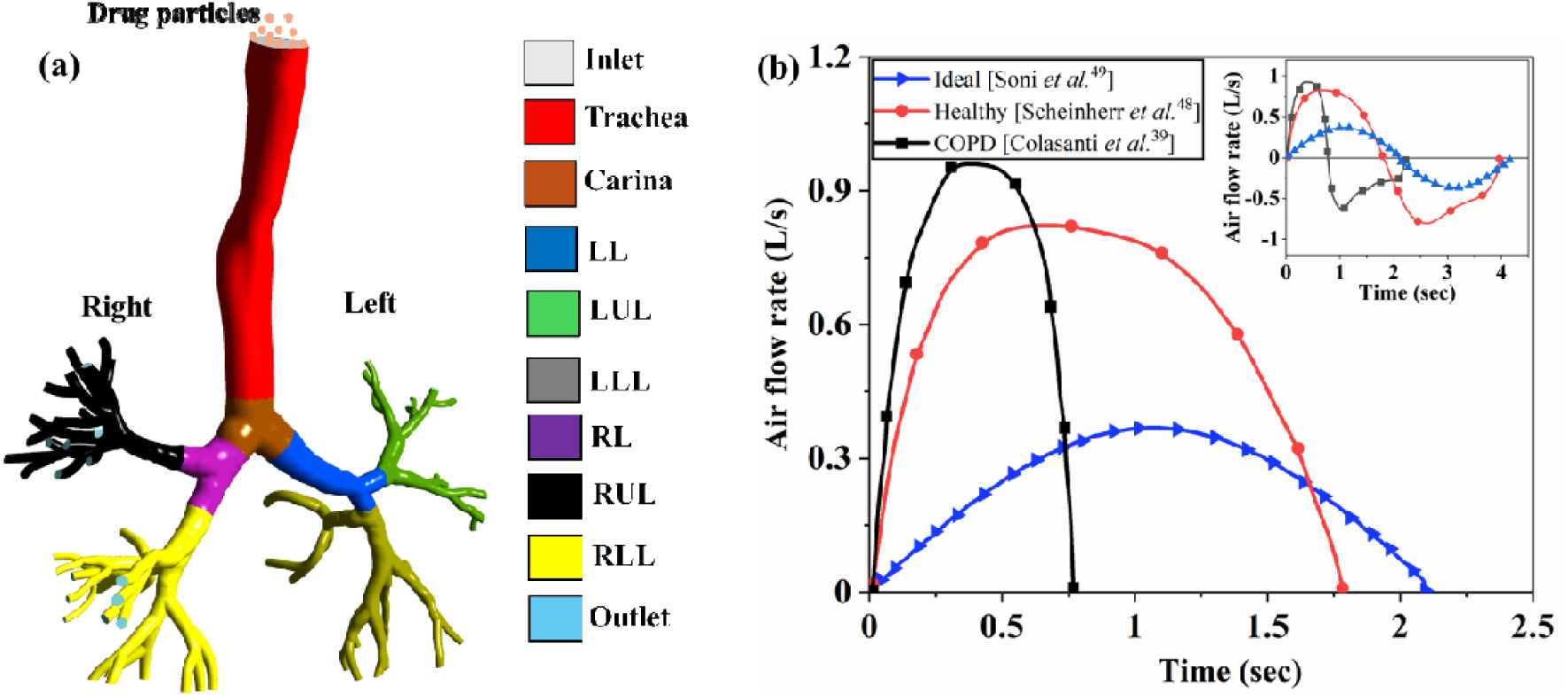
Lung airway model with different breathing profiles. (a) A CT-scan-based model of the bronchial airway up to G7 (Koullapis et al. 2018). The color tagging on this model illustrates different lobar segments and the inlet plane. (b) The comparison of airflow inhalation profiles for ideal, healthy, and COPD conditions, focusing on disease-induced differences in peak flow rate and breathing time. The inset plot shows complete inhalation-exhalation breathing cycles of the same.

A realistic breathing cycle was replicated by implementing a time-varying inlet condition (Fig. 2(b)). We employed the inhalation profile of around 50 COPD adult patients with severe airway occlusion, as described by Colasanti et al. (2004), to simulate compromised breathing. However, no artificial narrowing is applied to the airway segmentation because such obstructions usually occur downstream in the smaller airways (higher generations), which are not modeled in the current study and thus are not visible in the segmentations (Colasanti et al., 2004; Jeffery, 2004). The diseased inhalation profile of COPD shown in Fig. 2(b) closely resembles the profile described by Vitacca et al. (1999) for individuals with COPD recovering from an exacerbation. The peak inspiratory flow rate (PIFR) of the breathing waveform is 0.96 liters per second (L/s). A healthy breathing profile with a PIFR of 0.82 L/s was adopted from clinical data by Scheinherr et al. (2015) and used as the inlet condition for healthy cases. An idealized waveform is created from sinusoidal functions representing tidal breathing (Soni & Aliabadi, 2013). Based on this, we reconstructed the inhalation profiles of a 25-year-old individual with a peak flow rate of 0.36 L/s. Zero-gauge pressure is imposed at all outlets, and the walls of the airways are considered no-slip and stationary. Table 1 lists the parameters used in the computational simulation.

**Table 1.** Values of simulation parameters.

| Simulation parameters | Value | References |
| --- | --- | --- |
| Air density (kg/m<sup>3</sup>) | 1.225 | Bhardwaj et al. (2022) |
| Air dynamic viscosity (kg/m-s) | $1.7894 \times 10^{-5}$ | Bhardwaj et al. (2022) |
| Drug solution density (kg/m<sup>3</sup>) | 1000 | Bhardwaj et al. (2022) |
| Particle diameter (μm) | 1 to 10 | Ahookhosh et al. (2021) |
| Total number of injected particles<sup>#</sup> | 2,98,000 | - |
| Simulation time (sec)** | 2.14, 1.78, 0.78 | - |
| Time step size (sec) | 0.001 | - |

<sup>#</sup>2,98,000 of each particle size are injected at the trachea inlet.
\*\*The total simulation time for ideal, healthy, and COPD cases is 2.14, 1.78, and 0.78 seconds, respectively.

### 2.2. Airflow model

Since air is the working fluid in this study, it is assumed to behave as a Newtonian and incompressible fluid (Kim et al., 2019). The airflow dynamics inside the lung are characterized by the mass and momentum conservation equations (i.e., Navier-Stokes equations).

Mass conservation:

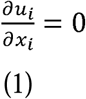

Momentum conservation:

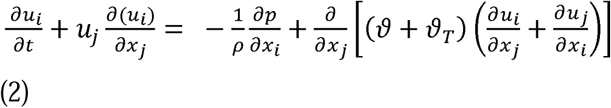

where, *ρ* denotes the density of the air, *ϑ* and *ϑ_T_* represent the coefficients of kinematic molecular, and turbulent viscosity, respectively. Within the lung model, airflow typically operates in a transitional turbulent regime, physically characterized by the presence of a wide range of time and length scales in the flow field. Furthermore, the inlet Reynolds number at the trachea is approximately 1900 for ideal breathing, 3242.3 for healthy breathing, and 5702.73 for COPD breathing. For such flow conditions, the k-ω SST turbulence model is selected for this study due to its proven effectiveness in capturing complex respiratory flows, particularly under wall-bounded, low Reynolds number conditions.(Jiyuan et al., 2015; Kleinstreuer & Zhang, 2010) This approach involves solving a hybrid transport equation for turbulent kinetic energy (*k*), dissipation (*∈*), and specific dissipation rate (*ω*),(Tiwari et al., 2021; Verma et al., 2024) along with the continuity and momentum equations. The transport equations for turbulent kinetic energy (*k*) and specific dissipation rate *(ω)* can be expressed by, *k* equation:

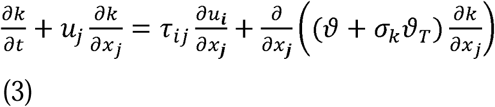

*ω* equation:

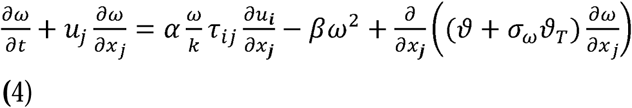

where, *τ_ij_* represent the viscous stress tensor. Further, the turbulent kinematic viscosity can be approximated empirically as 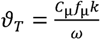, and the function *f*_μ_ represents as 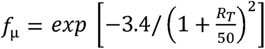 with 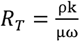. The additional constants in the above equations are defined as:

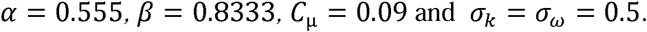

The pressure-velocity coupling is done by a SIMPLE algorithm, while the governing equations for momentum, turbulent kinetic energy, and specific dissipation rate are discretized using a second-order upwind scheme. The convergence criteria for all equations are set to 10^−6^ to ensure numerical accuracy and solution stability.

### 2.3. Particle transport

A one-way coupling is employed between the continuous (air medium) and the discrete (liquid particles) phases, and the volume concentrations of particle release are kept below 0.1% (Bhardwaj et al. 2022). Many studies have neglected particle-particle interactions, as the particle suspension becomes sufficiently diluted upon reaching the pulmonary airways (Rahman et al. 2022). Thus, the current model employs a one-way coupling approach, in which particle motion is driven by airflow dynamics while the effect of particles on the airflow is neglected (Kuga et al. 2023). A total of 298,000 spherically shaped particles are uniformly injected from the trachea inlet. In this study, we use the DPM-EWF to examine the interactions between continuous and discrete phases.

#### 2.3.1. Discrete phase model (DPM)

The Eulerian-Lagrangian DPM approach is implemented to simulate the trajectories of the particles within the lung airways. Particles are introduced from a surface upstream of the airway entrance to ensure sufficient dispersion before entering the trachea. Instead of solving a transport equation for them, each particle’s trajectory is computed individually in the Lagrangian frame of reference by solving its equation of motion, allowing for detailed tracking of particle behavior throughout the domain. The equation is stated as follows (Rahman et al., 2021b).

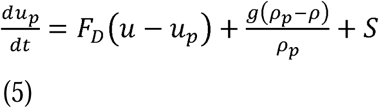

where, *F_D_*(*u* - *u_p_*) reflects the drag force per unit mass on the drug particles, and the last term accounts for the gravitational force.

The *F_D_* can be further expressed as,

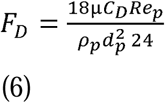

where, *u* indicates the velocity of air, *u_p_* signifies the drug particle velocity, *ρ* denotes the density of the air, μ is the dynamic viscosity of air, *ρ_p_* and *d_p_* symbolizes the density and diameter of the drug particle, respectively, and *C_D_* represents the drag coefficient.

Relative particle Reynolds number (*Re_p_*) is given by

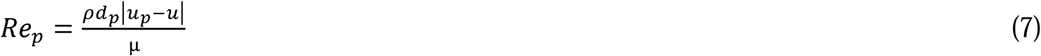

In Eq. (1), the variable *S* represents the additional forces that can influence particle motion under specific conditions. These forces are Saffman’s lift force, Brownian force, virtual mass force, and thermophoretic force. Thermophoretic force is neglected in this study due to the negligible temperature gradient. Similarly, the Brownian force is usually relevant in sub-micron-sized particles, and therefore, ignored for the current particle range (Ahookhosh et al., 2019; Arsalanloo et al., 2022; Kuga et al., 2023). The virtual mass force becomes significant when the carrier phase (air) density is higher than the discrete phase density 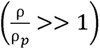. However, this is not the prevailing situation in the present investigation, rendering this force insignificant. Saffman’s lift force, caused by shear significantly affects sub-micron particles, is excluded here as the particle diameters ranges from 1 to 10 μm. Consequently, only drag and gravitational forces are considered significant in the present analysis (Arsalanloo et al. 2022). A stochastic tracking method was employed to capture the fluctuations in the turbulent flow field during simulation with the discrete random walk model and a random eddy lifetime. These components are defined under the turbulent dispersion parameters, which enable a more realistic modeling of particle trajectories inside the fluctuating turbulent flow field. Furthermore, particles reach the computational domain via the inlet surface. For particle deposition, a “trap” boundary condition is applied to the airway walls, while an “escape” boundary condition is applied at outlets (Ahookhosh et al. 2021). The escape condition allows particles to exit through the outlet without being reflected. Under this trap boundary situation, the tiny droplets adhere to the airway walls, forming a thin layer of drug solution coatings. The dynamics of these thin drug layers are further investigated using the EWF model.

#### 2.3.2. Eulerian wall film (EWF) model

The film model represents a two-dimensional thin liquid film on a wall surface. This thin film forms when liquid particles strike a solid surface within the domain. Upon impact, several outcomes are possible, such as the stick, rebound, spread, and splashing of the drug particles. Out of all these, the current study has considered the spreading and splashing of the droplets, which allows the formation of a thin liquid coating and the formation of secondary droplets upon impaction. The fundamental assumptions of the EWF model suggest that the film consistently flows parallel to the surface, which implies that the normal component of the film velocity is zero (Verma et al., 2024).

The governing equations of the EWF model are given as (Khao et al., 2023; Verma et al., 2025)

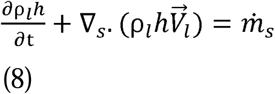

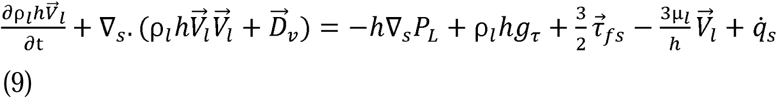

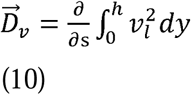

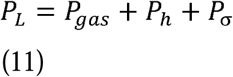

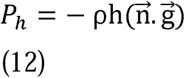

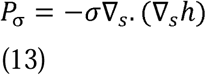

where, *ρ_l_* depicts the film density, *h* signifies the height of the film, *V_l_* represents the average film velocity and *ṁ_s_* indicates mass source per unit wall area resulting from particle deposition, film stripping, film separation, and phase change. *D_ν_* is the differential advection term and is calculated using the quadratic film velocity. *P_L_* designates the film pressure, *g_τ_* the gravity component, *τ_fs_* represents the shear stress at the liquid-film interface, *μ_l_* symbolize the viscosity, *q̇_s_* denotes the momentum source, *P_σ_* indicates the pressure contribution due to the surface tension, *P_h_* corresponds to the component of gravitational pressure acting perpendicular to the wall, and *n* represents the normal vector to the wall surface.

#### 2.3.3. Advantages of the EWF model over the interface capturing approach

The EWF model offers several advantages over other interface-capturing approaches in simulating thin liquid films in complex flow environments, such as those found in respiratory applications. Unlike the volume-of-fluids method (a popular interface capturing method), which requires a high mesh resolution to capture the liquid-gas interfaces and often results in higher computational costs, the EWF model simplifies the process by treating the film as a separate two-dimensional layer on the wall. This approach significantly reduces computational cost while still enabling detailed predictions of film dynamics, including film thickness, velocity, and droplet breakup. Additionally, EWF is well-suited for conditions involving large surface areas with thin liquid coatings. Its ability to couple with discrete phase models also enhances its capability to simulate droplet-wall interactions, making it an efficient and robust tool for applications requiring detailed wall film behavior with manageable computational expense.

#### 2.3.4. Scope of the study

Several limitations of the current investigation are acknowledged here, which indicate potential directions for future work to enhance the applicability of these findings in the context of inhalation therapy.

- The study focuses primarily on the inhalation phase, since reliable data on particle retention in deeper lung areas are unavailable (Koullapis et al., 2019; Koullapis et al., 2020). Furthermore, there is no defined system for tracking particles that remain airborne and re-enter during exhalation, which limits the precision of mimicking the entire breathing cycle.
- Another simplification involves the assumption of rigid airway walls. In reality, the airway tissues exhibit some degree of wall elasticity, which can influence airflow and deposition patterns.
- The current study ignores mucus layer formation to simplify the computational setup. The mucus layer under diseased states (COPD and asthma) exhibits stiff viscoelastic behavior, which allows it to behave as a soft solid wall (Abrami et al., 2024; Lai et al., 2009; Leal et al., 2017). Under such conditions, droplet impaction may eventually lead to thin film formations and their spread over the surface. Moreover, the explicit modelling of the viscoelastic mucus layer, along with air and droplets, will potentially create a three-phase system, which is currently ignored and reserved for future investigations, considering its reduced effects on the thin film spread dynamics under diseased conditions.
- The study assumes a constant droplet size, which simplifies modeling and reduces computational costs. While evaporation can influence droplet dynamics, it has a secondary effect under the controlled ambient conditions and short inhalation times. This assumption allows us to focus on core deposition and film generation mechanisms without overcomplicating the model.

Despite these limitations, the simulation results for airflow behavior and particle deposition align well with findings in existing literature, lending confidence to the overall validity of the study.

### 2.4. Computational approach

A flowchart (Fig. 3) depicts the process of setting up and solving a coupled DPM-EWF simulation in ANSYS Fluent 2023 R1. The main steps involved are as follows:

**Fig. 3.**
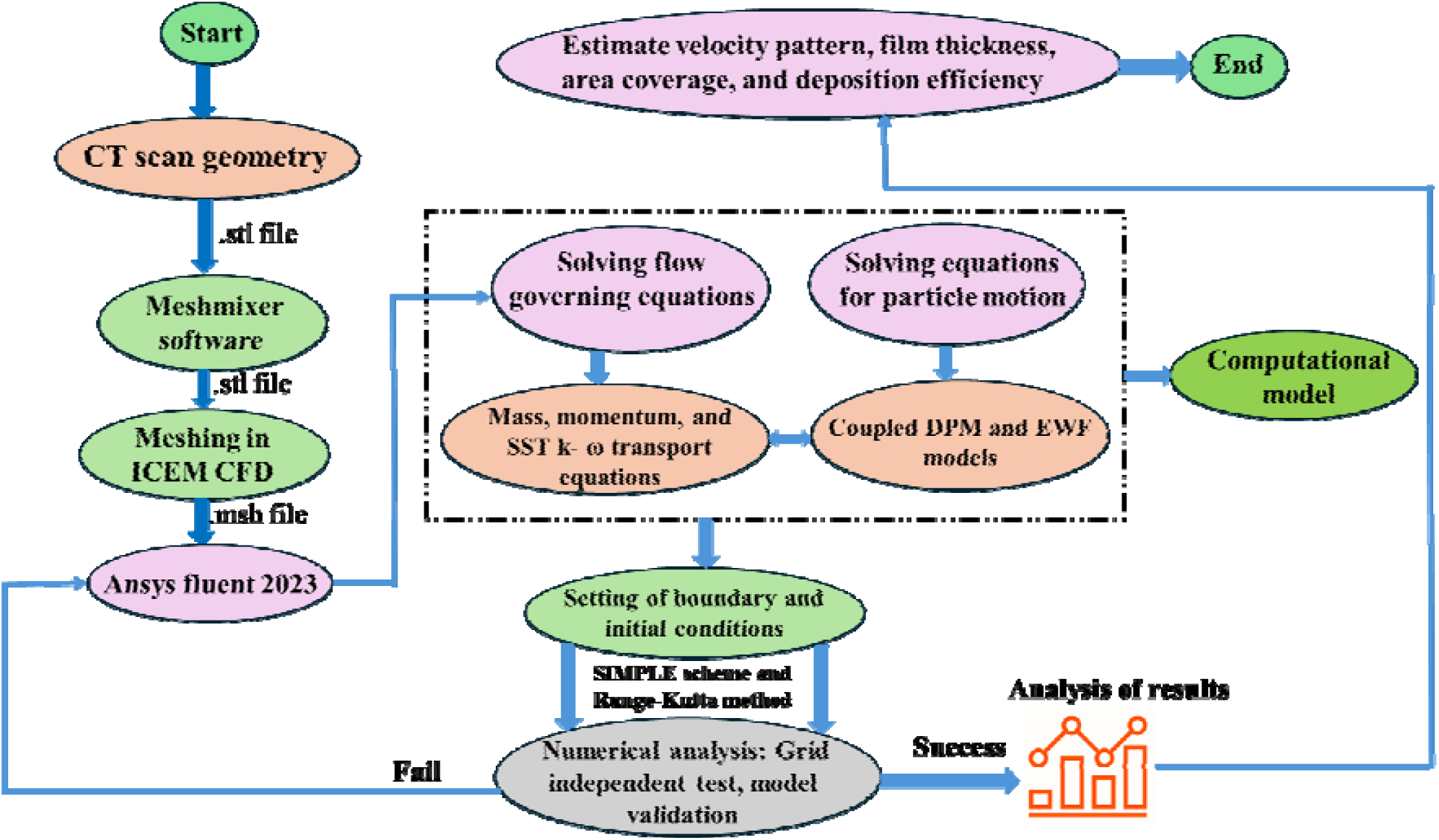
The flowchart of inhaler-based liquid drug particle delivery simulations in a CT-scan-based lung model. The procedure starts with geometry construction and meshing to coupled airflow-particle modeling using DPM and EWF methods, followed by validation and performance evaluations.

First, the CT scan geometry is smoothed, and artifacts are removed using Meshmixer software developed by Autodesk^®^ to ensure a clean and accurate representation of the airway structure. The computational domain is then imported into ICEM CFD in *.stl* file format to create a high-quality computational grid. After creating the mesh, the *.msh* file is imported into ANSYS Fluent, where the boundary conditions and fluid flow properties are employed and configured. Following that, initial conditions and particle properties, such as diameter, density, and discrete phase settings, are specified. Once the initial settings have been established, a suitable turbulence model is chosen depending on the flow parameters. The governing equations for fluid flow (Eqs. 1–4), particle motion (Eqs. 5–7), and thin film dynamics (Eqs. 8–13) in the coupled DPM-EWF model are then numerically solved, taking into account appropriate boundary conditions for both the fluid and particle phases. Before initiating the primary flow study, the complete computational setup is thoroughly tested to ensure adequate grid quality. This procedure includes evaluating the results against experimental and computational benchmarks. Section E and F contains more information about the grid independence study and validation techniques. The entire computational investigation was performed on a Dual AMD Genoa 9th series high-performance computing (HPC) processor (128 cores) with 1 TB RAM. The overall computing time to simulate the process for one model is around 74 hours.

### 2.5. Grid dependency test

The computational grid (Fig. 4) is generated in ICEM CFD, which consists of unstructured elements with 10 layers of near-wall prism to adequately capture flow dynamics near the boundary more precisely, as seen in Fig. 4(a). A grid dependence test was performed to ensure the mesh density was sufficient to achieve converged results. The meshing procedure employs robust octree and quick Delaunay techniques to improve mesh quality and computational performance. The mesh dependency study included three meshes with element counts ranging from 3 million (3M) to 9M (Fig. 5). The analysis of velocity magnitude fluctuations along the two locations of the human lung revealed negligible changes (0.027%) when the mesh density was increased from 6 to 9M elements. As a result, the current study used a mesh density of 6M elements for the rest of the results.

**Fig. 4.**
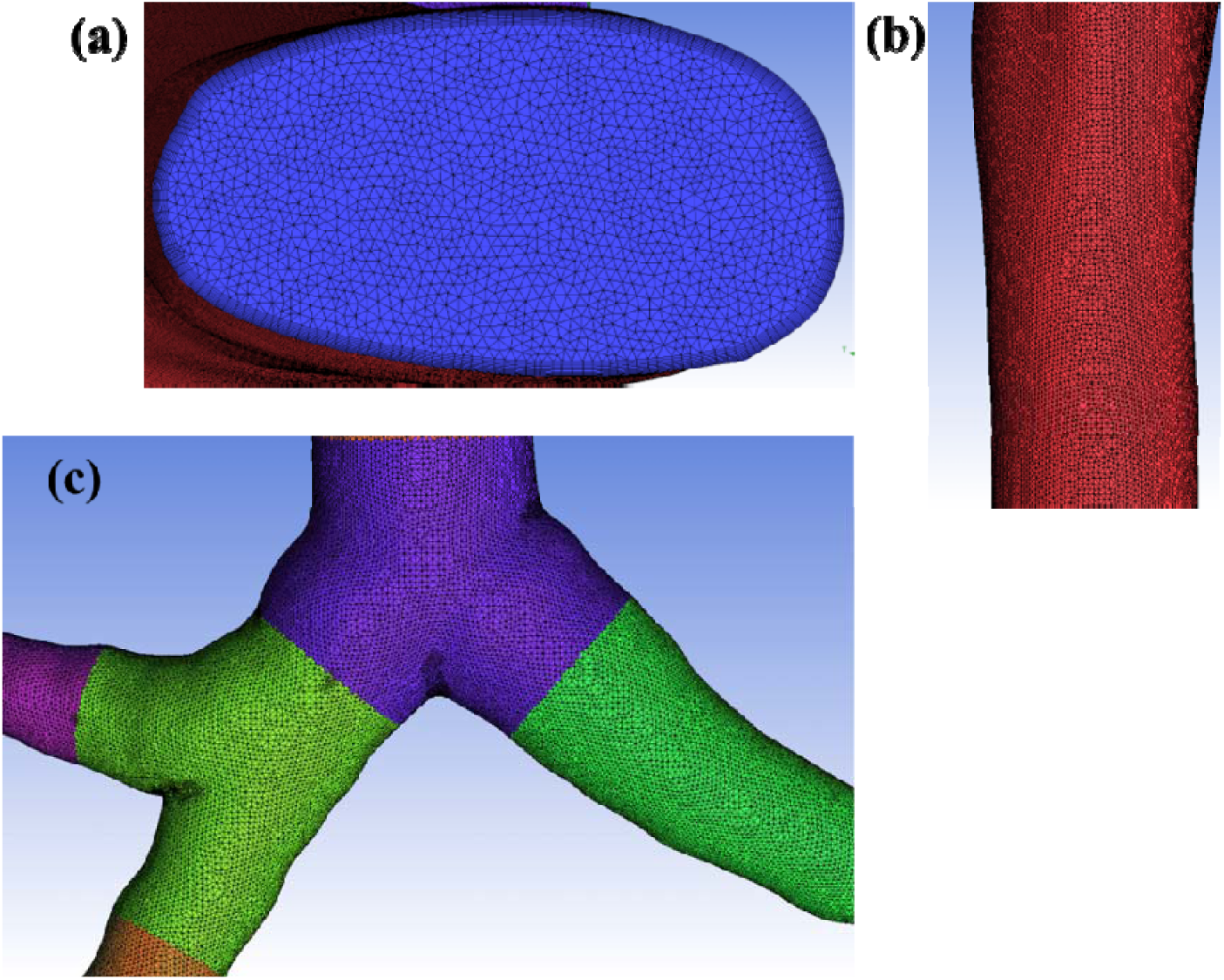
Schematic representation of meshing strategy for a complex, realistic geometry of respiratory airways along the (a) inlet plane, (b) tracheal, and (c) bifurcating regions. The figure highlights the unstructured tetrahedral grid distributions with 10 inflation layers along the airway wall.

**Fig. 5.**
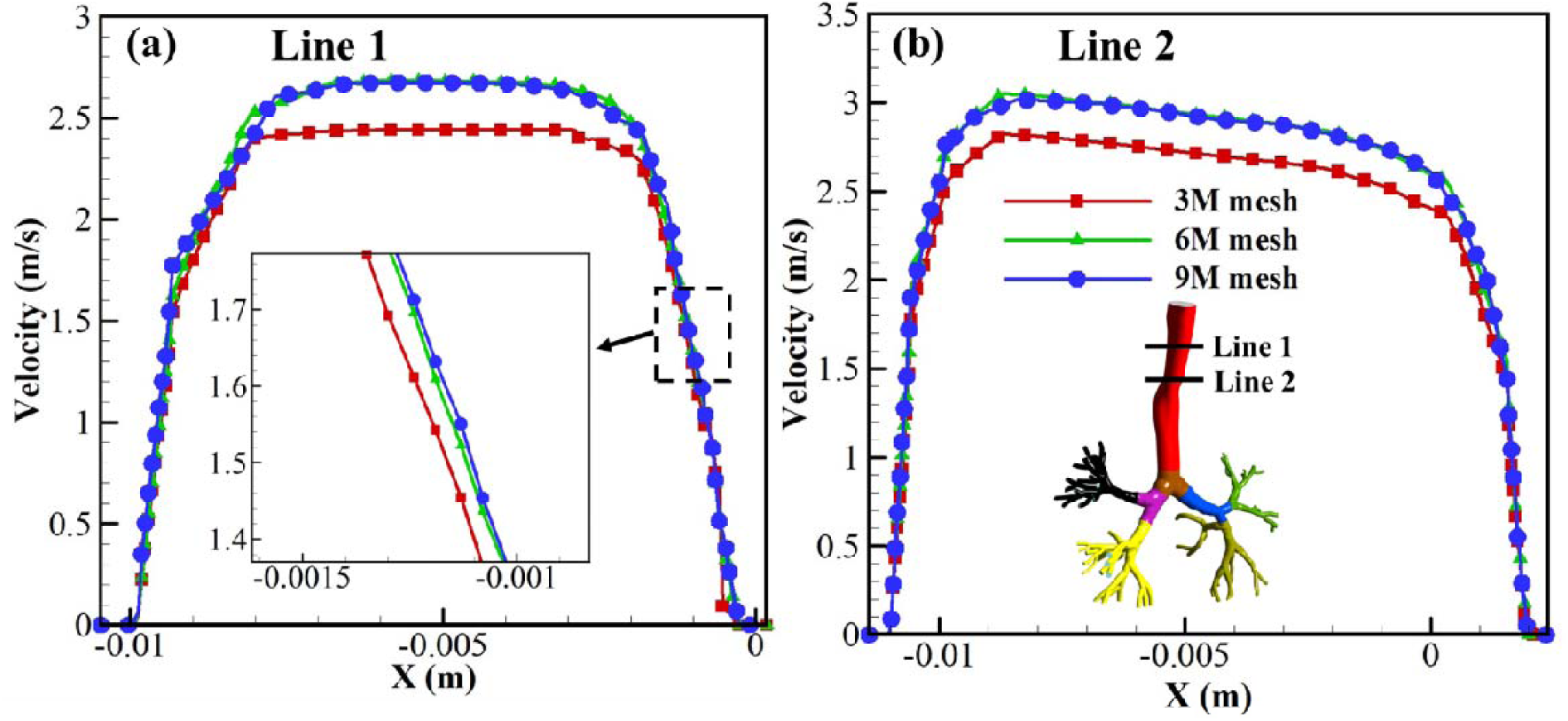
Grid independence test compares velocity distributions obtained using different mesh resolutions along (a) line 1 and (b) line 2. The results demonstrate negligible changes in the velocity profile between 6M and 9M meshes. Thus, the ‘6M mesh’ is chosen as an optimal grid for the rest of the simulations.

### 2.6. Model Validation

The credibility of the present computational framework has been firmly established through systematic validation efforts reported in our prior studies. The DPM-EWF model was previously established on a Weibel’s geometry by Verma et al. (2024), where the validations were shown against both experimental (Kim & Fisher, 1999) and numerical (Kim et al., 2019) results. Such elaborate validations demand a distinct simplified geometry instead of the realistic airway presented in the current paper. However, the base simulation setup in both cases is identical. Therefore, to avoid redundancy, the corresponding validation plots in our previous work (Verma et al., 2024) are referred here rather than presented explicitly. Additionally, the geometric accuracy of the airway model used in the current investigations has been verified by comparing the anatomical reconstructions reported by Bhardwaj et al. (2022) demonstrating good agreement in terms of structural fidelity and flow behavior. Combinedly, the validated model setup and geometrical fidelity provide confidence in the accuracy and reliability of the current results.

## 3. Results and discussion

This study examines the deposition efficiency of drug particles under three distinct breathing conditions: ideal, healthy, and diseased (COPD). The particle sizes ranging from 1-10 μm (Ahookhosh et al., 2021) are investigated, assuming a perfectly spherical shape throughout their transport. This section analyzes airflow dynamics and particle behavior inside the human respiratory airways.

### 3.1. Airflow dynamics

Fig. 6 presents the flow characteristics across different regions of the respiratory tract. Fig. 6(a) shows the transverse cross-section of the trachea, serving as a reference for subsequent lobe-wise analysis. These planes extend from the trachea and cover various levels of branches and bifurcations within the left and right lungs. Fig. 6(b) illustrates the velocity distribution and flow streamlines for three different breathing rates: ideal, healthy, and COPD. The peak inlet velocity for an ideal, healthy, and COPD case is around 2.1, 4.9, and 10.1 m/s, respectively. It has been observed that even minimal geometrical deflections can modify the streamlines’ uniformity, resulting in the formation of secondary vortices. This can be seen in plane B, where a small vortex forms (more visible in the COPD case) at the upper side of the plane as a result of deflection caused by the asymmetrical bulging of the tracheal cross-sections. Secondary vortices are also detected further downstream due to abrupt changes in the flow path, irregular bifurcations, and various deflections in the airway geometry. The flow divides into two parts at the first bifurcation, but because the left main stem bronchus is slightly narrower than the right, the flow accelerates accordingly. This is marked on planes C (left) and D (right). Although the geometry remains constant for all three cases, COPD patients frequently inhale more forcefully due to greater airway resistance in real-life situations, resulting in a PIFR. The high-velocity flow in the simulation mimics the real-world scenario in which COPD patients try to compensate for diminished lung efficiency by breathing faster and harder. The finding indicates that, even in an anatomically normal airway, changing breathing profiles can considerably impact airflow behavior and potentially affect particle deposition, oxygen transport efficiency, and overall pulmonary function.

**Fig. 6.**
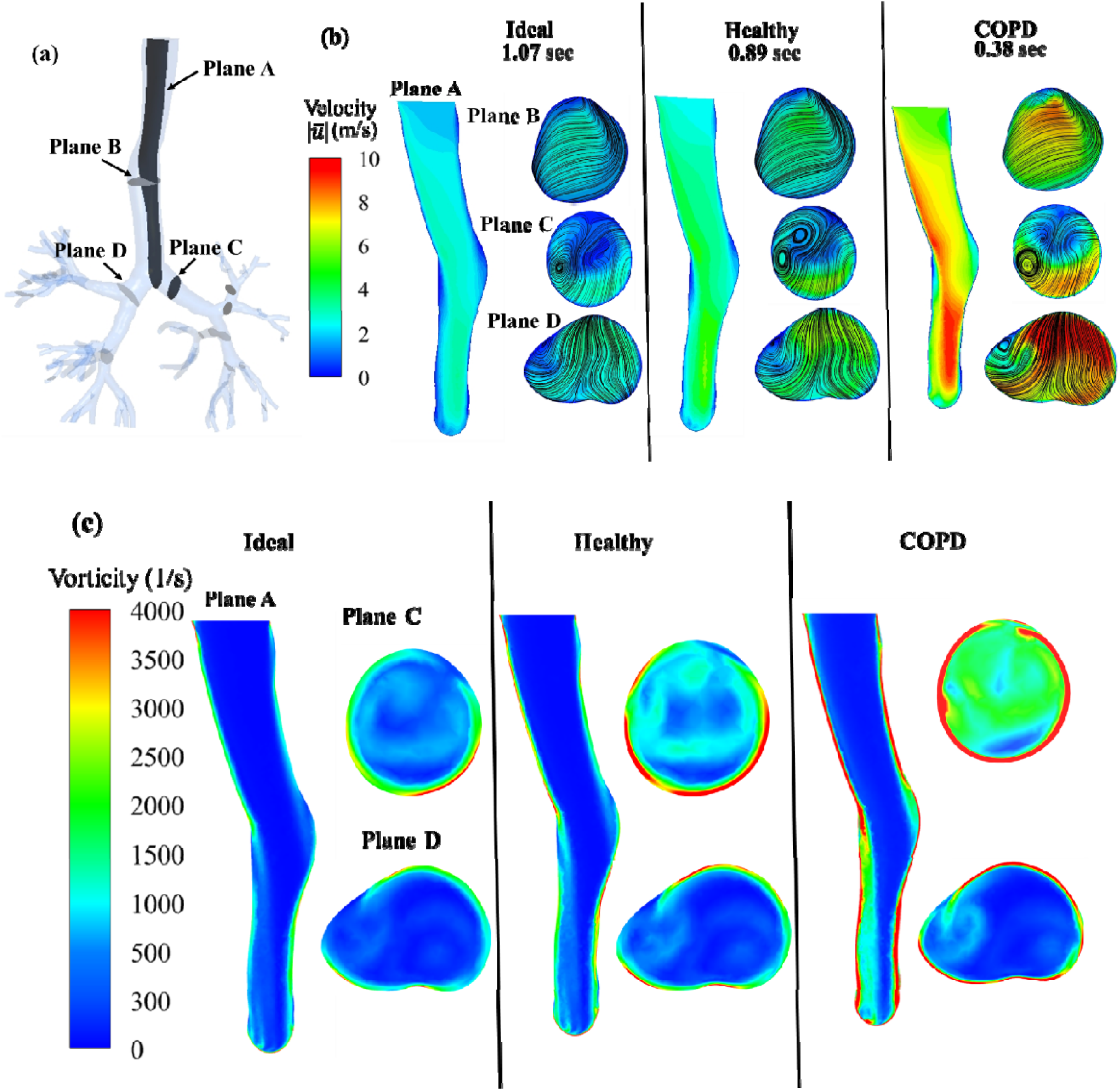
Plots representing (a) the location of the cut-sectional planes at which the airflow parameters are investigated, (b) velocity distributions, and (c) vorticity magnitude contours at different cut-planes of the human airway. The contours are extracted at the PIFR of all three ideal, healthy, and COPD breathing profiles.

The comparison of the change in vorticity magnitudes (Fig. 6(c)) highlights the dominating complex rotational flow field of the COPD case compared to the other two. Vorticity is maximum at the outer walls rather than at the center due to the presence of strong velocity gradients and shear layers, which resembles that of Bhardwaj et al. (2022). As can be seen, the maximum vorticity contrasted from 2400 s^−1^ (ideal breathing) to 4000 s^−1^ for the COPD case. In general, plane C has higher vorticity than plane D in all three cases. The COPD breathing depicts the highest vorticity (~4000 s^−1^ in Plane C, ~2500 s^−1^ in Plane D), followed by healthy (3500 s^−1^ in Plane C, ~2500 s^−1^ in Plane D), and then the ideal breathing (~3000 s^−1^ in Plane C, ~2000 s^−1^ in Plane D). The no-slip boundary condition and high shear near the wall cause vorticity gradients towards the center locations. At airway bifurcations, flow redirection causes secondary motions and swirling structures (such as Dean vortices), contributing to increased vorticity near the boundaries. In conditions such as COPD, the turbulent fluctuations become more prominent, and the high shear leads to the maximum turbulent kinetic energy values of 0.4 m^2^/s^2^ compared to ideal and healthy values of 0.007 and 0.015 m^2^/s^2^, respectively. Furthermore, recirculating flow zones near bifurcations produce additional rotating effects, increasing vorticity in these areas. In contrast, the center portion of the airway has a more stable and uniform flow with little velocity fluctuations, resulting in less vorticity. The combination of velocity gradients, boundary layer effects, turbulence, and secondary flow structures results in maximum vorticity near the outer walls.

### 3.2. Deposition and escape pattern of the particle

Deposition efficiency is expressed as the ratio of particles trapped (absorbed) on the airway walls to particles entering the computational domain through the inlet. It is represented by *η_f_*, and can be expressed as,

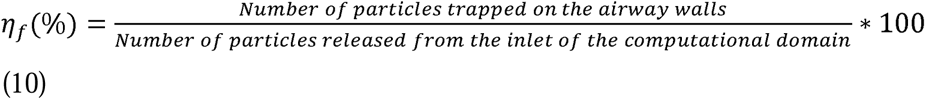

Fig. 7 depicts the regime map for the deposition and escape efficiency of particles of varying diameters under ideal, healthy, and COPD breathing conditions. The figure indicates that 10 μm particles have the highest deposition, and 1 μm particles have the least deposition throughout the domain, regardless of breathing rate. This is primarily because inertial impaction becomes less effective as particle diameter decreases. A similar trend has also been reported in a previous study by Gorji et al. (2015). In ideal and healthy cases, the LLL bronchus and LL exhibit the highest deposition rate for most particle sizes, while the carina, RL, and trachea have the lowest. However, in the case of COPD, deposition is highest at the carina, whereas the LLL, RLL, and trachea show the lowest deposition rates. Similar to the particle diameter, it has been observed that the increase in inhalation rate also monotonically increases the deposition efficiency and subsequently reduces the escape rate. This behavior can be explained by inertial impaction, which occurs when particles with high inertia are unable to follow the curved flow streamlines, especially at bifurcations or sharp turns in the airway.

**Fig. 7.**
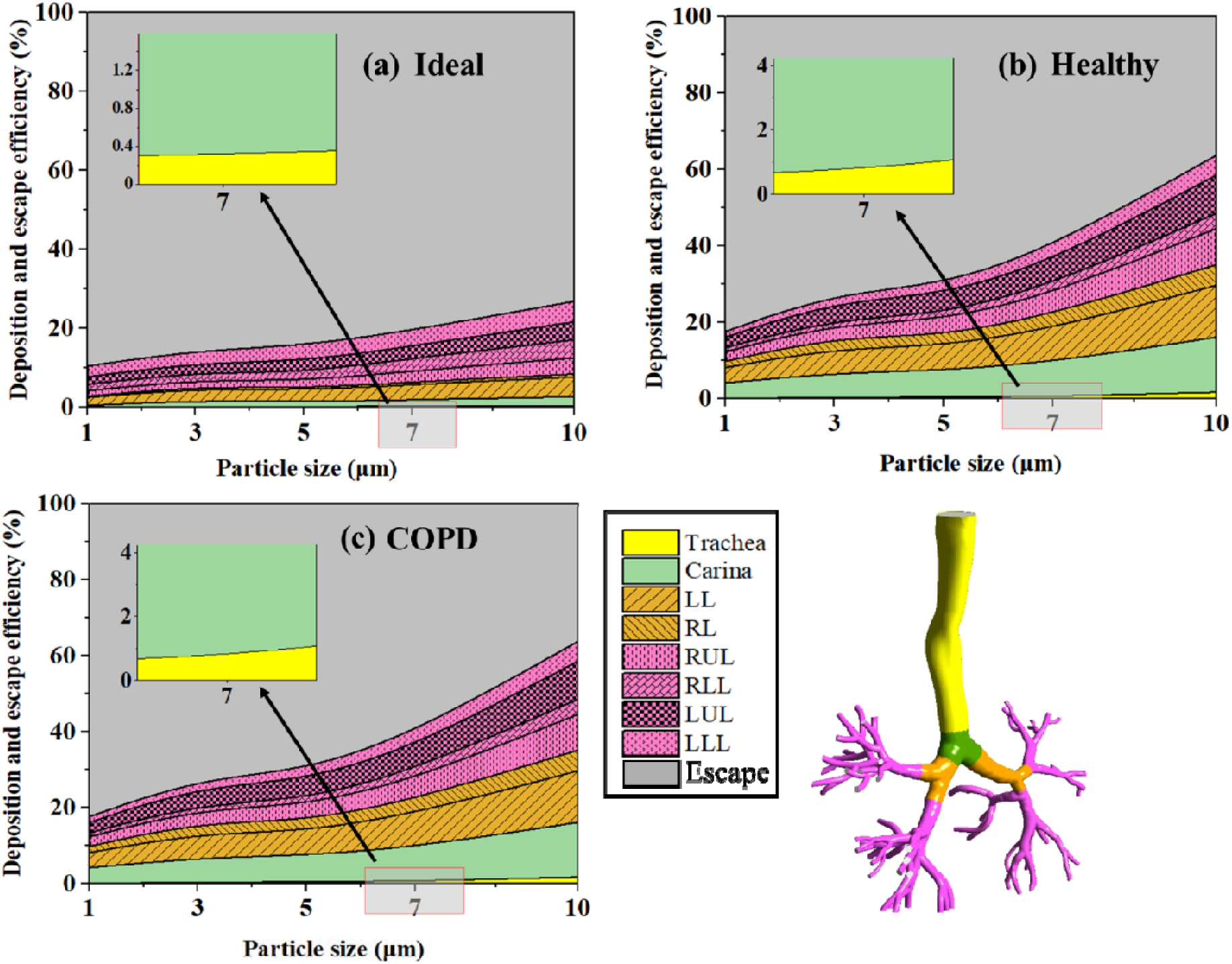
Regime maps showing the deposition efficiencies of inhaled particles across different lung regions and final escape percentage (grey color) from the entire domain till G7. Three distinct maps represent (a) ideal, (b) healthy, and (c) COPD breathing conditions. The deposition patterns are grouped based on different colored lung regions, as illustrated in the last image.

Changes in flow direction amplify this effect, causing larger particles to deviate from the airflow and impact the airway walls. Consequently, most of the large particles deposit in the upper airway generations, and penetrations to the deeper regions of the lungs are reduced. Smaller particles (1-3 μm) are deposited primarily through diffusion, resulting in a consistent distribution throughout lobes. As particle size exceeds 5 μm, inertial impaction becomes the primary mechanism, resulting in a more distinct variation in deposition between upper and lower lobes. In the case of COPD, intake becomes more forceful and less effective as the airflow velocity changes abruptly, resulting in higher inertial impaction at bifurcations, particularly along the carina region. This limits the deep lung penetration in COPD cases, i.e., fewer particles reach the lower lobes. Moreover, the other complex dynamics, such as air trapping or the formation of local air pockets, make upper airway deposition more prominent in COPD cases.

Fig. 8 presents a comparative analysis of the total deposition efficiency of inhaled particles, sized between 1 and 10 µm, across the different breathing profiles, namely the ideal, healthy, and COPD cases. As explained earlier, the deposition efficiency of the smaller particles (1-3 µm) is relatively low for all three cases due to their high diffusivity and lower inertia. However, due to disturbed airflow patterns and high velocity, the deposition is still relatively higher in COPD cases (≈ 22.69%) compared to healthy (≈ 16.76%) and ideal (≈ 12.45%) breathing. For medium-sized particles (5-7 µm), the deposition efficiency of the COPD case starts dominating (nearly 34.77%) compared to healthy and ideal breathing by roughly 22.69% and 17.57%, respectively. This rise in deposition efficiency can be attributed to a combination of inertial impaction and altered flow characteristics in COPD. For larger particles (10 µm), there is a significant difference in deposition efficiencies amongst the breathing profiles. It reaches up to approximately 1.2 and 2.37 times higher for COPD cases compared to healthy and ideal breathing, respectively. Conclusively, the short breathing span, formation of local air pockets, higher inertial impaction, and gravitational settling lead to less penetration of the liquid particles in COPD cases.

**Fig. 8.**
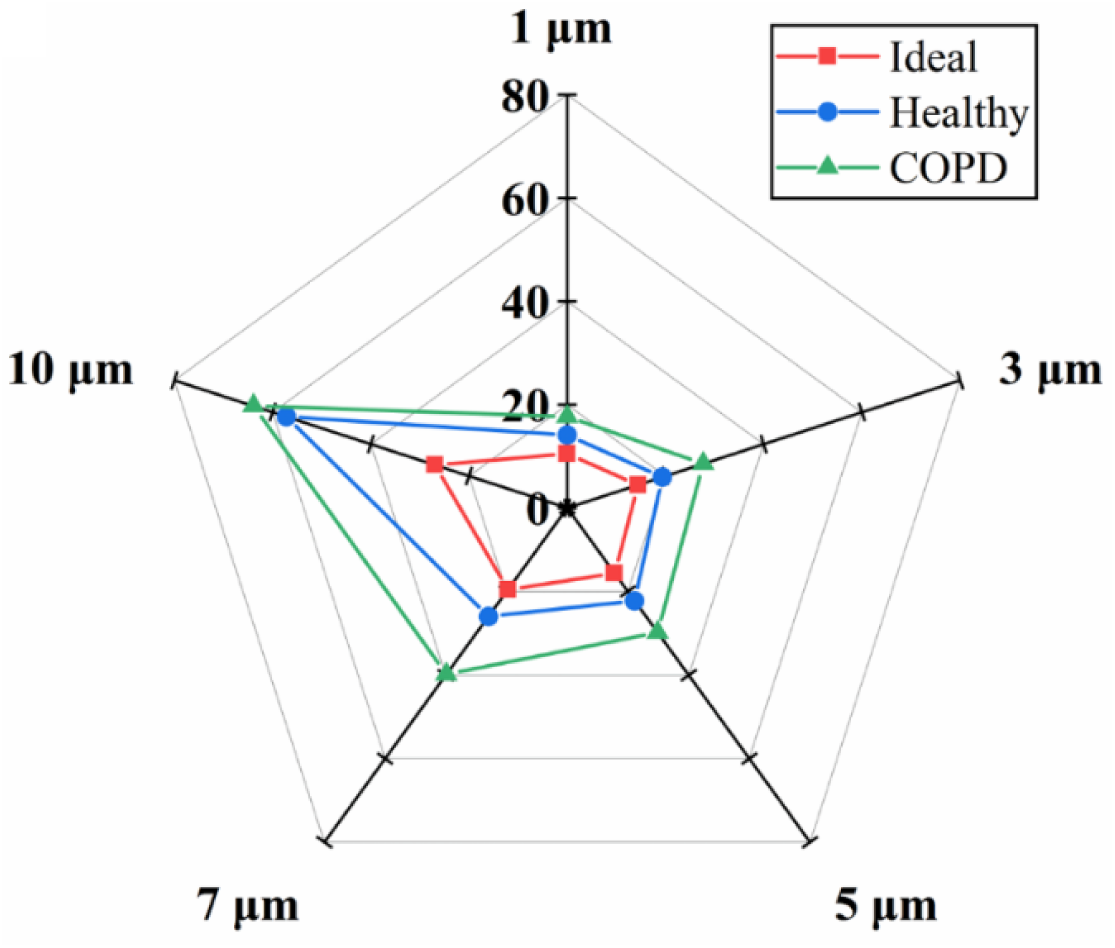
Total deposition efficiency across distinct particles for ideal, healthy, and COPD, highlighting the influence of breathing conditions on deposition behavior for each particle size.

Figure 9(a) shows that the left lung consistently has a higher deposition efficiency (roughly two times) than the right lung for all three cases of breathing. In an ideal breathing situation, the left lung has approximately 6.64% deposition efficiency, whereas the right lung has around 3.3%. In healthy breathing, deposition in the left lung increases significantly to about 8.21%, while the right lung remains below 4.5%. Although deposition efficiency in the left lung increases marginally in COPD patients (8.99%) compared to healthy individuals, it remains higher than in the right lung (5.15%). The primary reason behind the deposition bias is a considerable disparity in airflow distribution between the two lungs. Specifically, the average velocity magnitude in the right lung is around 38% lower than in the left. This difference in velocity magnitude is because of an early bifurcation (middle image in Fig. 9(a)) along the right bronchus in the airway geometry. This leads to flow stagnation immediately after the first bifurcation and directs a greater proportion of the flow to the left bronchus. Moreover, an asymmetry in the bifurcation angle at the carina also plays a decisive role in the preferential biases of the airflow rate. As a result, fewer particles enter the right lung, whereas increased velocity toward the left lung encourages more particle transport and deposition in that region.

**Fig. 9.**
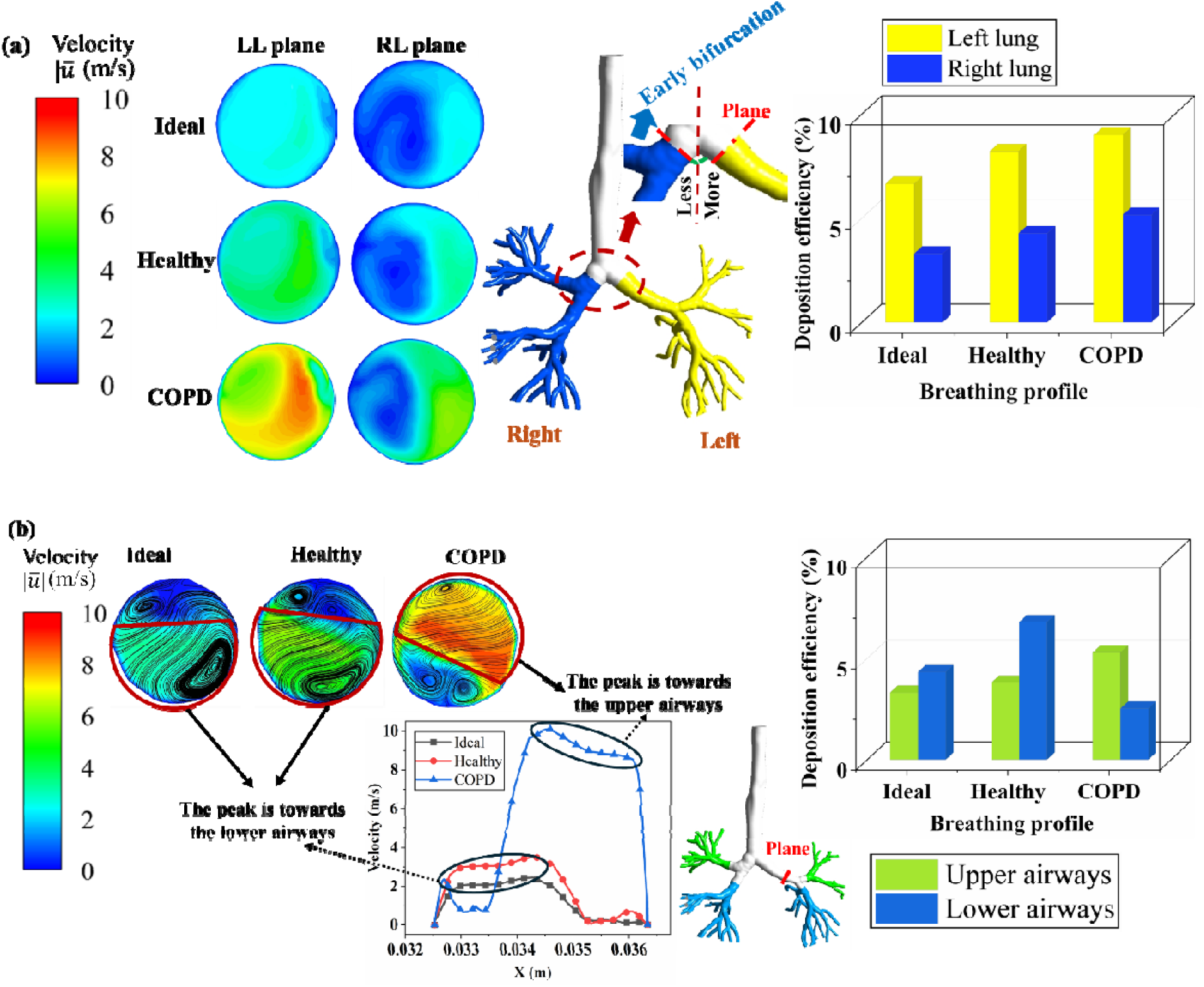
Comparison of particle deposition efficiency in various lung regions for 1 μm particle size. (a) Particle deposition asymmetry between left and right lungs (right), which potentially arises because of the early bifurcation at the right lung (middle) and differences in average velocity magnitude (left) as a result of it and (b) deposition efficiency contrasts between the upper and lower lung lobes (right) that arise because of the preferential asymmetries of velocity magnitudes, highlighted using the contour (left) and line (middle) plots.

Fig. 9(b) highlights the preferential biases in deposition efficiency between the upper and lower lung branching. As shown in the bar plot of Fig. 9(b), the deposition efficiency in the lower airways is higher than in the upper airways for both ideal and healthy breathing profiles. However, this trend is completely opposite for the COPD breathing profile. Here, the upper airways display higher deposition (~5.28%) than the lower airways (~2.52%).

The complex airflow dynamics are the fundamental reason behind this preferential asymmetry in deposition efficiency. It has been known and widely established that every airway branching introduces an asymmetry in the velocity distributions (Liu et al., 2023; Verma et al., 2024). However, in the current situation, this asymmetry in the flow velocity is tweaked in such a way that forced and shortened breathing of COPD leads to a shifting of the velocity peak to the upper side of the airway (left images of Fig. 9(b)). As a result, more particles are directed towards the upper airway branches. Meanwhile, in both ideal and healthy breathing circumstances, peak velocity is predominantly centered in the lower portion of the airway. This directed momentum drives more particles into the lower airway branches, increasing deposition in those areas. The overall understanding of deposition preference suggests that COPD has different deposition profiles than normal and idealistic breathing, on which most inhalers are usually designed. Therefore, a disease-specific inhaler-based drug delivery demands special attention to achieve therapeutic outcomes.

Fig. 10 depicts a symbolic representation of particle distributions across airway generations. The 1 μm particles scatter more evenly throughout the lung regions than the 5 and 10 μm particles. Smaller particles have lower inertial impaction, allowing them to closely follow fluid streamlines and lessen their chances of colliding with airway walls. As a result, these particles can escape the computational domain and travel into the deeper lungs. Larger liquid particles, on the other hand, are more likely to collide with the airway wall, resulting in more deposition in targeted lung sites.

**Fig. 10.**
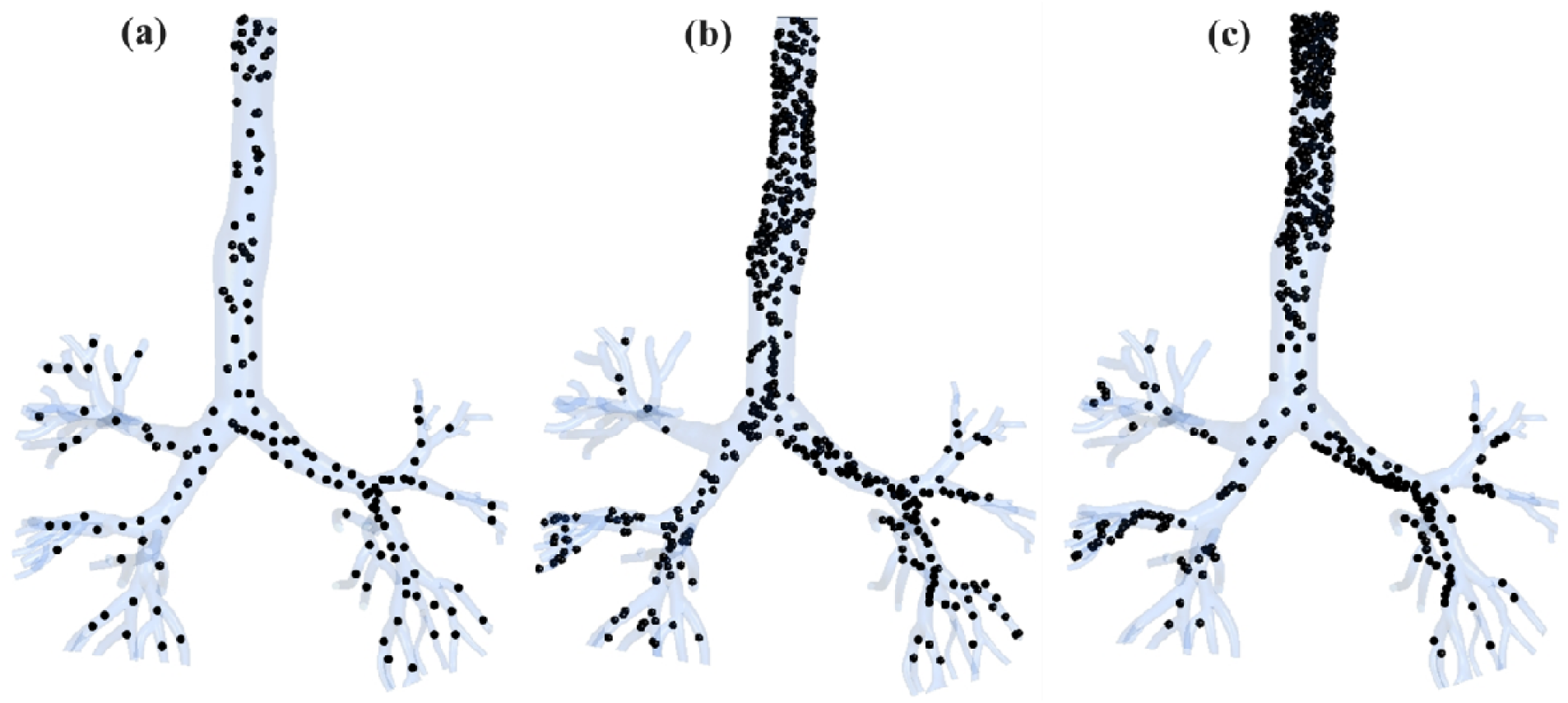
Symbolic representations of spatial particle distribution to highlight the effect of inertial impaction by the (a) 1 μm, (b) 5 μm, and (c) 10 μm particle sizes under healthy breathing conditions.

### 3.3. Film thickness

The film thickness is the height of the thin liquid layer formed on the airway walls by the deposition and coalescence of tiny droplets. This deposition, combined with existing liquid layers, forms a thin coating. The variation in film thickness depends on different factors, such as the particle shape and size, airway geometry, breathing pattern, and inherent flow dynamics. Figs. 11(a) and 11(b) show typical film thickness distributions for the normal breathing profile and 10 μm particle size. The asymmetry in the velocity distributions at different locations of airways is evident in the contour plots of velocity magnitudes (Fig. 6(b)). This asymmetry leads to selective deposition with each generation (Fig. 11(a)). As can be seen, over 90% of particles are deposited along the inner segment of the airway walls rather than the outward regions. To justify the role of velocity asymmetry, a plot of velocity variations along the diameter is shown on the right bronchi [Fig. 11(b)]. The asymmetry in deposition is more pronounced for 5-10 μm particles than for smaller ones [Fig. 11(c)].

**Fig. 11.**
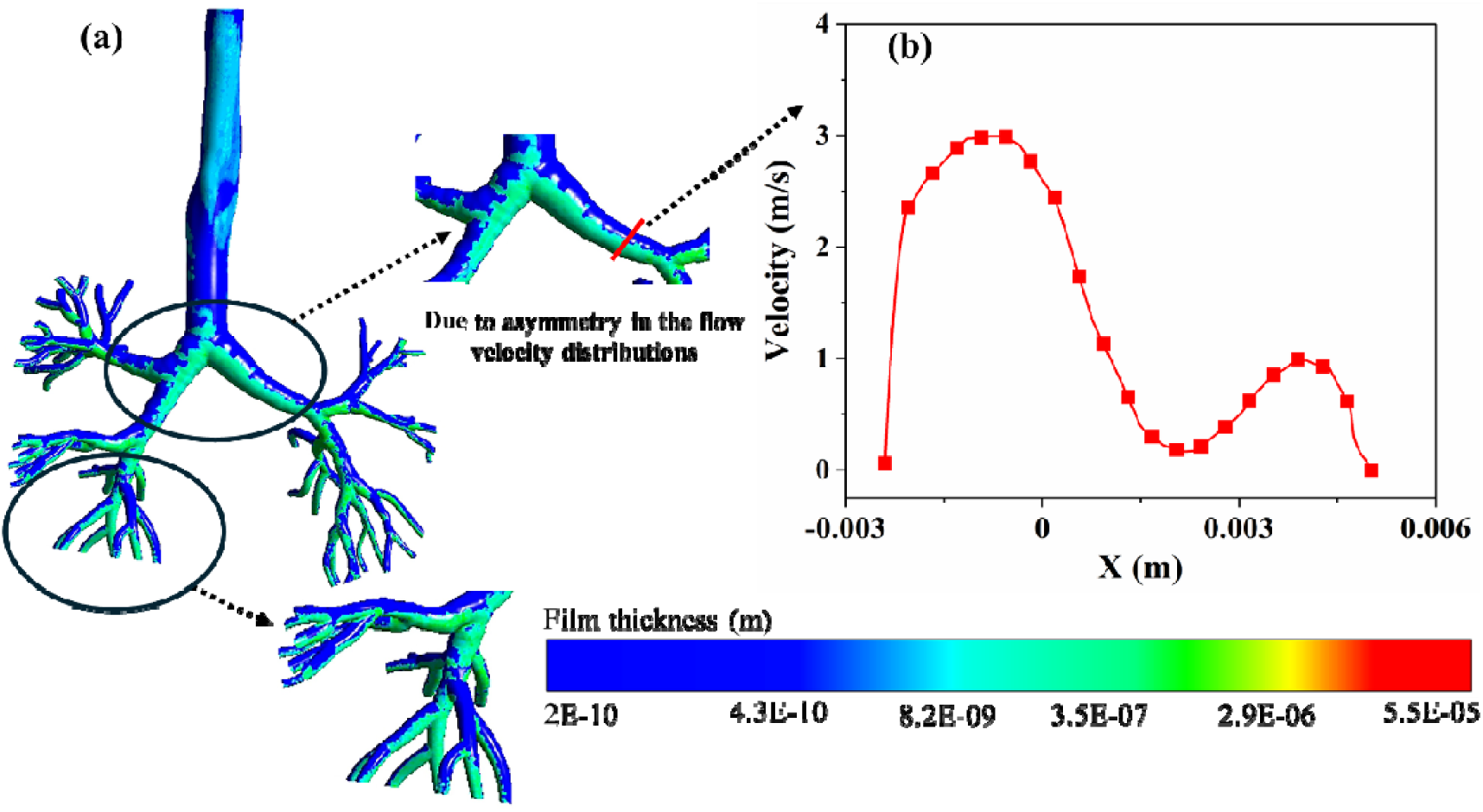
(a) Contour plots of film thickness to represent the particle depositions along inner segments of airways, and (b) the asymmetric variations in the velocity magnitudes across the diameter of the airway cause such non-uniformity of the film thickness.

**Fig. 11(c).**
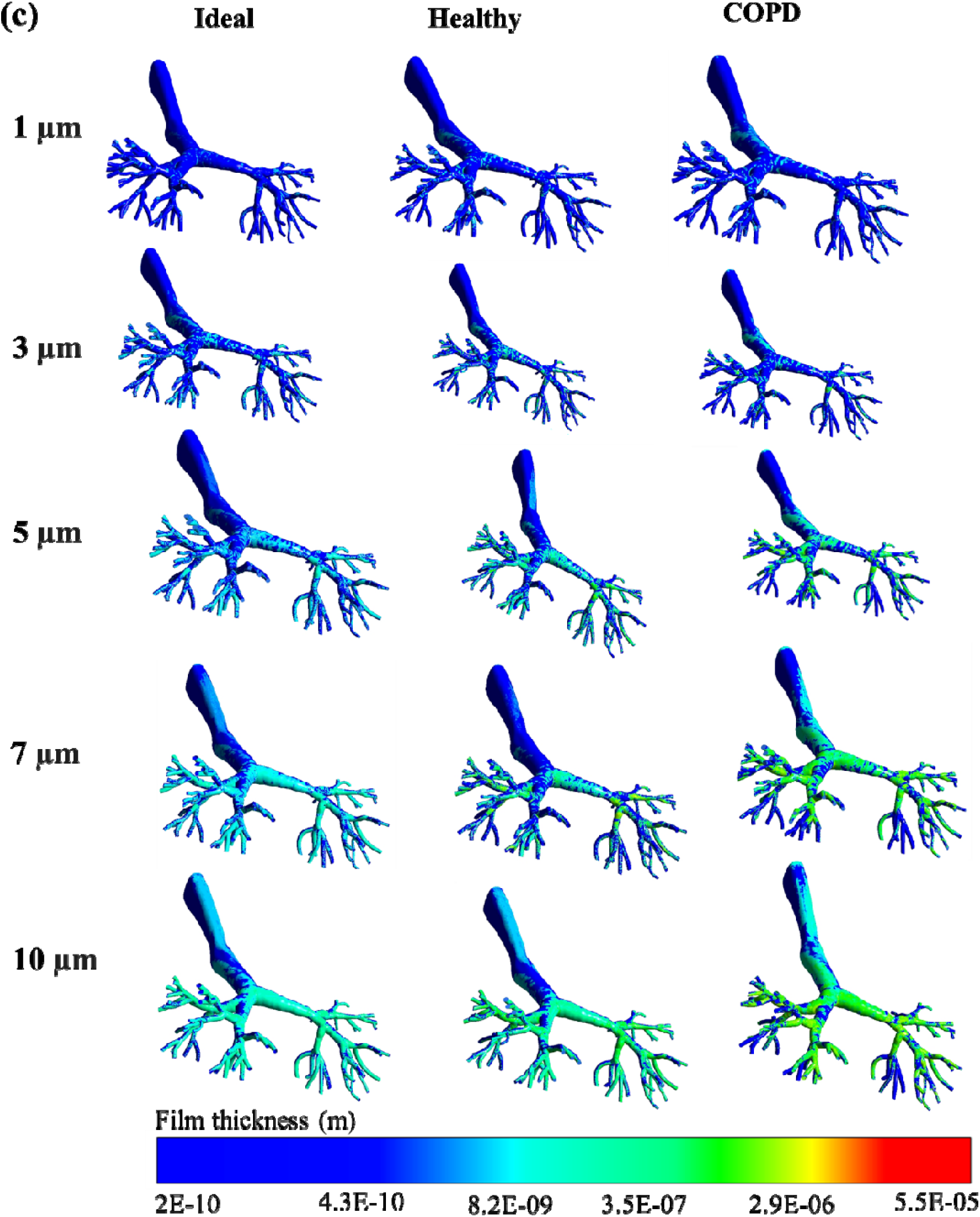
Comparison of film thickness variations for the particle sizes ranging from 1 to 10 µm under all three inhalation cases. Higher thickness appears for the larger particle size due to inertial impaction, whereas the forceful breathing of the COPD patient leads to a relatively higher film thickness.

The film thickness plots highlight the importance of resolving the post-impaction spread dynamics of the inhaler-based drug delivery mechanism. It captures the precise location of particulate deposition and overall area coverage to give a comprehensive picture of drug delivery efficacy. Generally, the zonal subdivision terminology of airways defines the entire annular wall with 360° rotations as a single zone. As a result, the previously employed pure DPM approach cannot capture the porosity in depositions along any designated airway zone. In contrast, the EWF allows us to capture not only the precise location of depositions but also the post-striking dynamics, such as the spreads and interactions of initially deposited drugs caused by the shear of the surrounding airflow. Thus, relying on just DPM-based approaches compromises the accuracy of drug delivery predictions.

A similar trend of change in deposition efficiency with the particle size and different breathing patterns is evident in the variation of film thickness comparison shown in Fig. 11(c). However, the film thickness distribution on these plots suggests that the overall extent of area coverage in each zone has significant porosity. In the case of COPD breathing, film thickness is more prominent in the upper airways compared to both ideal and healthy breathing due to the characteristics of altered airflow dynamics. COPD patients experience increased airway resistance and turbulence, which promote impaction and particle deposition in the upper bronchial region. The constrained airflow reduces particle delivery to the lower airways, producing less film accumulation. This situation significantly impacts drug delivery efficacy because of the complex airflow dynamics that prevent effective deep lung deposition.

Compared to the healthy case, the ideal breathing has the smallest film thickness for all particle sizes. The natural variations in airflow rate, acceleration, and turbulence in healthy breathing enhance inertial impaction and convective transport of particles. In contrast, the ideal breathing profile shows a smooth, idealized inhalation cycle without the strong acceleration peaks that are found in normal human breathing patterns. The absence of considerable turbulence (turbulent kinetic energy ≈ 0.007 m^2^/s^2^ for the ideal and 0.015 m^2^/s^2^ for the healthy) minimizes impaction, resulting in less deposition in the airway. Instead, the particles diffuse uniformly throughout the airways, resulting in a thinner film layer in ideal breathing. Medically, sinusoidal breathing does not entirely resemble physiological conditions, but it does provide a simple model for demonstrating how controlled airflow reduces impaction.

Fig. 12 compares how the film thickness at different locations changes with the change in particle size for all three breathing cases. The distribution of maximum film thickness at different zones follows a similar trend in both the ideal and healthy breathing situations (Figs. 12(a) and 12(b)). The left lung consistently has the maximum film thickness, which is nearly 2.63 times greater in the ideal case and roughly 5.94 times greater in the healthy case than the right lung thickness for all the particle sizes in an averaged sense. However, the film thickness in the trachea is almost negligible and significantly lower in the carina (15 times for healthy and 23.7 times for ideal breathing) compared to the maximum left lung depositions in the healthy and ideal breathing situations. In contrast, the COPD instance shows the completely opposite trend for maximum film distributions (Fig. 12(c)). Unlike the other two, the maximum film thicknesses are observed in the carina region rather than the left or right lungs. The maximum film thickness is roughly 1.48 times lower for the left and 2.83 times lower in the right lung compared to the maximum carina region. Moreover, it has been observed that the general distributions of maximum film thickness in almost all the regions for COPD cases are nearly an order of magnitude higher (10^−6^ m) than in ideal and healthy breathing situations. The shortened and forceful breathing in COPD cases introduced a contrasting difference in the distribution and overall deposition of maximum film thickness. This suggests that the change in the breathing pattern significantly influences the inhalation airflow dynamics inside the airways. This leads to the formation of flow recirculation, secondary vortices, higher turbulent intensity, and increased airway resistance.

**Fig. 12.**
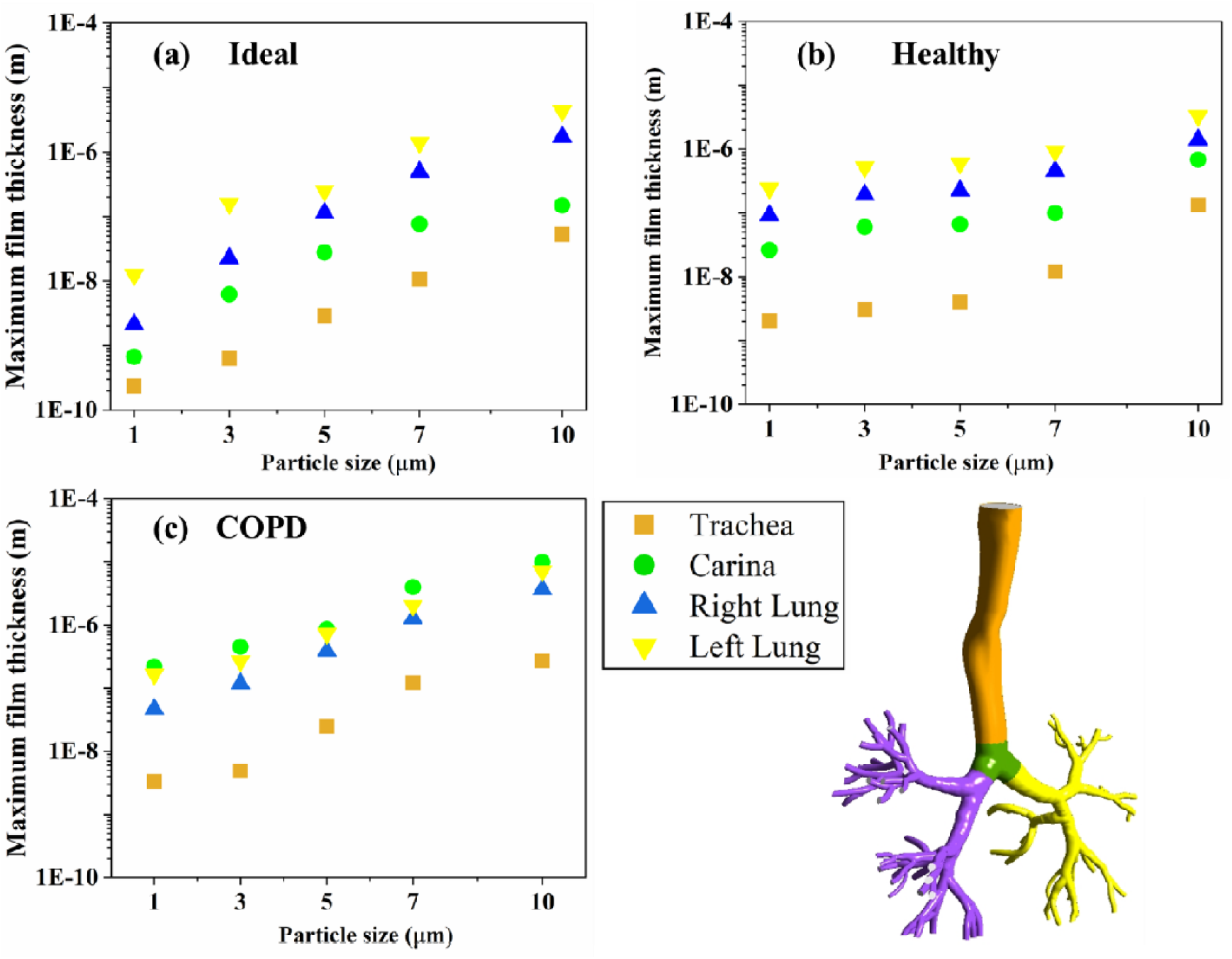
Comparison of film thickness deposition for various particle sizes (1-10 µm) in the different airway segments under (a) ideal, (b) healthy, and (c) COPD conditions. The Y-axis displays maximum film thickness variations on a logarithmic scale. The color tagging of the symbols is linked with the equivalent zone for better understanding (bottom right image).

### 3.4. Film area coverage

The film area coverage (Fig. 13) is the percentage of a surface covered by a liquid film after deposition, expressed as the wet area fraction. In other words, it is the proportion of the entire surface area covered in each individual lobe by a liquid film generated by drug solutions upon impact. This parameter is critical in inhalation drug administration since it influences how well the medication spreads along the airway lining. Smaller particles are less impacted by inertia and can remain suspended in the airflow for longer periods, minimizing their tendency to settle and deposit on the airway walls. The area coverage plot is critical in understanding the spatial pattern of particle deposition throughout the individual regions of the respiratory tract. Here, capturing area coverage is made possible by coupling the thin film approximation (EWF) approach with the DPM-based particle tracking. In contrast to deposition efficiency, which measures the total fraction of inhaled particles that deposit anywhere in the lungs, area coverage focuses on where those particles settle and what fraction of the airway surface area is covered.

**Fig. 13.**
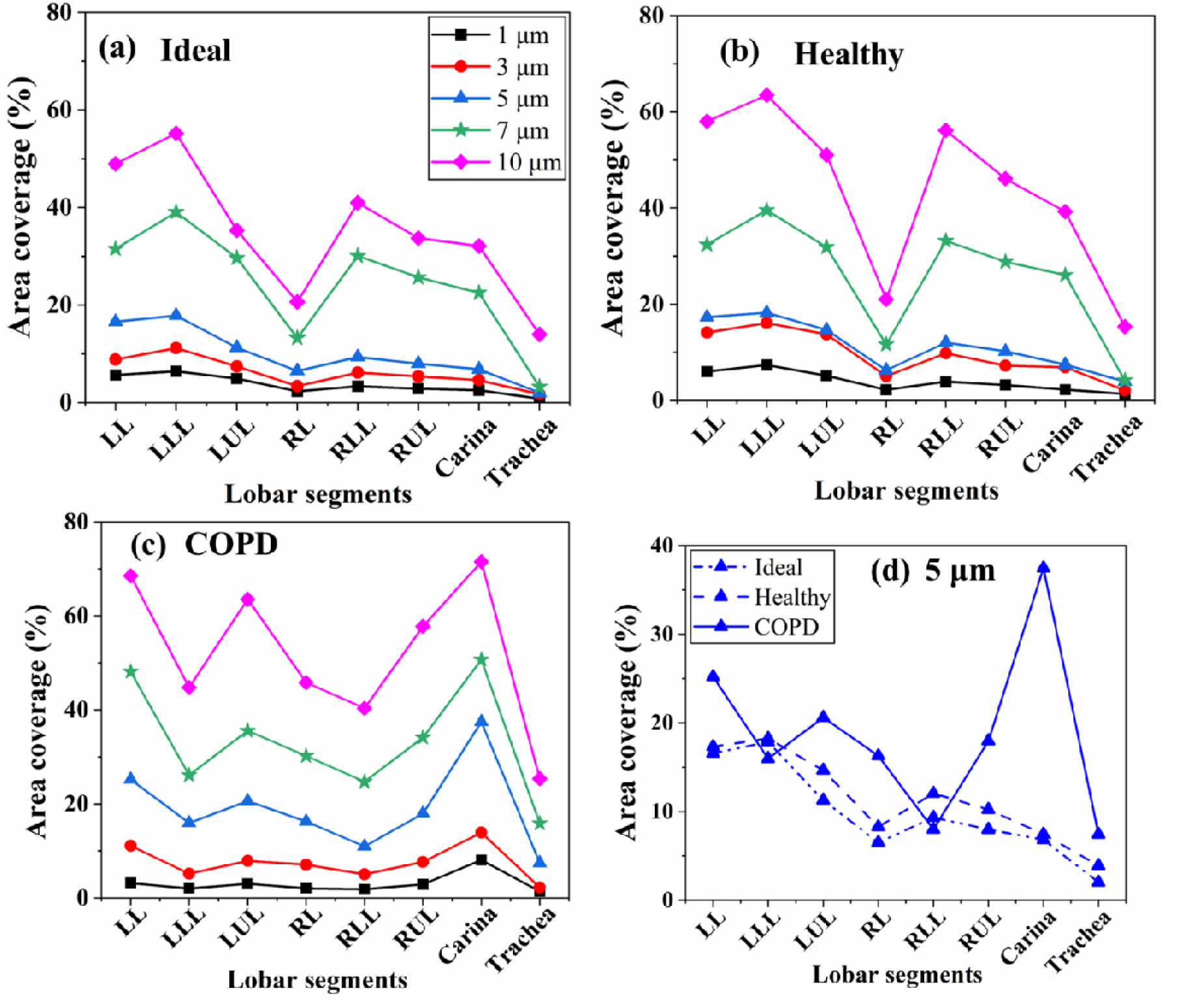
The variations in area coverage (%) along the different lobar sections with particle sizes ranging from 1-10 µm for the (a) ideal, (b) healthy, (c) COPD cases, and (d) comparison of all three breathing profiles for 5-µm particles.

In Fig. 13, the four major regions mentioned initially, namely the trachea, carina, right, and left lungs, are further segmented into eight different zones to predict the deposition behavior more precisely. Specifically, the right and left lung regions are further split into the RUL, LUL, RLL, and LLL lobes. This segmentation enables us to understand the greater resolution of spread patterns to identify region-specific accumulation. Such denotations are helpful in the clinical assessments of targeted drug delivery. The distributions reveal that the LLL region shows the highest area coverage for both ideal (55.2%) and healthy (63.48%) breathing profiles (Fig. 13(a) and 13(b)), followed by the LL, LUL, and the right-side airways (RLL, RUL, and RL). The inlet breathing profile of COPD completely changes this distribution. The carina exhibits the maximum area coverage at about 71.51% for the same 10 µm particle size, followed by the upper (LUL, RUL) and lower (LLL, RLL) lobar segments. Such region-specific coverage provides essential insights about the drug delivery and its spread efficacy, which would otherwise be impossible to calculate when only the DPM-based approach is used in drug delivery investigations. The area coverage spread variations with the particle size show a more or less similar pattern of distributions. Therefore, breathing profiles-wise comparison for only one particle size (5 µm) is shown in Fig. 13(d). Each breathing state has a specific pattern in which the drug spreads across the airway surfaces. Under COPD breathing, some lobar segments have much larger area coverage, most notably the carina, which peaks at around 38%, followed by LUL and RUL. This reflects the impact of the rapid and forceful inhalation characteristics of COPD, which redirect the airflow and concentrate deposition in upper bifurcation zones. In contrast, healthy and ideal breathing profiles have more consistent but lower film coverage across the lobes, with no sudden spikes. The healthy profile has relatively higher area coverage than the ideal one, implying that natural inhaling dynamics result in more extensive distribution than simplified flow profiles. Overall, the findings suggest that disease-specific breathing dramatically modifies surface coverage patterns, which should be modeled effectively for better disease-specific drug delivery efficacy.

## 4. Conclusions

This study utilizes a CT scan-based realistic human lung model, extending up to the G7 from the trachea (G0-G7), to analyze the efficacy of particle depositions. A numerical investigation examines the effects of five distinct particle sizes between 1-10 µm under three breathing condition rates (ideal, healthy, and COPD). Additionally, the coupled DPM and EWF model is employed to evaluate the deposition behavior of particle interactions within the respiratory airways.

The key findings of the current research work are outlined below:

- The velocity and vorticity distribution contours suggest the presence of a highly disturbed, rotational flow field because of the short and forceful breathing of a COPD patient compared to the ideal and healthy person’s breathing pattern. A variation in maximum vorticity magnitude is 4000 s^−1^ for a COPD patient compared to ideal (3000 s^−1^) and healthy (3500 s^−1^) breathing. Such a disturbed flow field significantly alters the deposition dynamics of inhaled drug particles and affects their distributions.
- The deposition efficiency has been observed to increase monotonically with the increase in particle size for all three transient breathings. This pattern varies from 12.45 - 26.96% for ideal breathing, 16.76% - 57.17% for healthy breathing, and 22.69% - 63.89% for the COPD cases in an averaged sense as the particle sizes are increased from 1-10 µm.
- In general, the short and forceful breathings of COPD have higher deposition efficiency than the other two cases for all respective particle sizes. In an average sense, this increase is roughly 2.05 and 1.28 times compared to ideal and healthy cases.
- COPD breathing shows higher deposition along the upper lung lobes, which otherwise have a contrasting preference in ideal and healthy breathing cases. Ideal and healthy breathing profiles show higher deposition along the lower lungs..
- The highest area coverage and maximum film thickness for the COPD is found in the carina, whereas, for ideal and healthy breathing, the same has been changed to the LLL.
- Thus, this study reveals that impaired breathing can significantly reduce the drug penetration into the deep lung compared to ideal and healthy inhalation. This reduction occurs due to altered airflow dynamics, increased airway resistance, and turbulence, which result in greater deposition in the upper airways and less drug reaching the lower respiratory regions.
- Therefore, advanced inhalers such as spacers, valved holding chambers with pMDIs, vibrating mesh nebulizers, and some breathing exercises are essential to slow particle velocity, allowing deeper penetration for better treatment. The study emphasizes the need to combine disease-specific modulation with dynamic inhalation patterns to enhance therapeutic efficacy.

## Acknowledgements

The authors are grateful for having the opportunity to use the high-performance computing (HPC) system at the Department of Mechanical Engineering, NIT Rourkela, through the ISRO-funded project “Design of Micro Cryogenic Coolers for Phased Array Receiver” (Project ID: RES-ISTRAC-2022-008). The author also acknowledges the funding from ANRF-PAIR (Project ID: SR-25-CR-047), title “Consortium of Technologies for Sustainable Agriculture, Health, Energy and Environment”, for meeting consumable needs.

## Author declarations

### Author contributions

**Sameer Kumar Verma:** Data curation (equal); Formal analysis (equal); Investigation (equal); Methodology (equal); Software (equal); Validation (equal); Visualization (equal); Writing – original draft (equal). **Saurabh Bhardwaj:** Conceptualization (equal); Formal analysis (equal); Project administration (equal); Supervision (equal); Writing – review & editing (equal). **Kishore Singh Patel:** Conceptualization (equal); Funding acquisition (equal); Project administration (equal); Resources (equal); Software (equal); Supervision (equal); Writing – review & editing (equal). **B. Kiran Naik:** Conceptualization (equal); Funding acquisition (equal); Project administration (equal); Software (equal); Supervision (equal); Writing – review & editing (equal).

### Data availability

The data that support the findings of this study are available from the corresponding author upon reasonable request.

